# An endosome-associated actin network involved in directed apical plasma membrane growth

**DOI:** 10.1101/2021.01.07.425786

**Authors:** LD Rios-Barrera, M Leptin

## Abstract

The subcellular membrane-trafficking system plays many roles in cell growth and morphogenesis, from bulk membrane provision to targeted delivery of membrane proteins and other cargos. In tracheal terminal cells of the Drosophila respiratory system, endocytosis and trafficking through late endosomes balances membrane delivery between the basal plasma membrane and the apical membrane that forms a subcellular tube, but it has been unclear how the direction of growth of the subcellular tube with the overall growth of the cell is coordinated. We show here that endosomes are involved in this process as coordinators of F-actin. Actin assembles around late endocytic vesicles in the growth cone of the cell, reaching from the tip of the subcellular tube to the leading filopodia of the basal membrane. Preventing actin nucleation at late endosomes disturbs the directionality of tube growth, uncoupling it from the direction of cell elongation. Severing actin in this area affects tube integrity. Our findings show a new role for late endosomes in directing cell morphogenesis by organizing actin, in addition to their known role in membrane and protein trafficking.

## Introduction

Cell shape changes require coordination between the actin cytoskeleton which determines shape, and the plasma membrane which must be delivered to and removed from the appropriate domains to allow the formation of new structures. Therefore, membrane-cytoskeleton interactions are important in shape transitions, since they transfer the forces generated by the cytoskeleton to the membrane at the cell surface, which will ultimately define the new shape (Sitarska and Diz-Muñoz, 2020). Shape changes also require subcellular regulation; migrating neural crest cells extend filopodia in the direction of migration to increase traction, while forming actomyosin contractions in the rear, propelling the cell forward (Piacentino et al., 2020). Cells undergoing apical constriction restrict an actomyosin network to the apical cell cortex and retrieve apical plasma membrane, while keeping a more constant shape in the basal domain of the cell (Gilmour et al., 2017). Studying how these mechanisms are controlled at the subcellular level in physiological contexts is often challenging, and little information exists on how they are coordinated to generate new cell shapes. Failure to establish proper cytoskeletal-membrane interactions and therefore proper cell shape is detrimental for development and tissue functionality, and it is also a major cause of human disease (Alekhina et al., 2017; Kasza et al., 2019; Shcherbata et al., 2007; Walton et al., 2016), highlighting the importance of studying these mechanisms.

Terminal cells of the Drosophila respiratory system are a useful model to study how cells execute cell shape changes from a subcellular perspective. These cells lie at the tip of the tracheal branches that allow gas exchange throughout the body of the animal. To reach the furthest positions, terminal cells form complex ramifications each containing an interconnected subcellular tube. This tube forms by the invagination of the apical plasma membrane, which successively continues to grow inwards while the cell elongates and branches. This leads to a complex cell morphology consisting of hollow tubes (Best, 2019; Hayashi and Kondo, 2018). The mechanisms that terminal cells use to build a subcellular tube resemble those in other lumen-building structures (Sigurbjörnsdóttir et al., 2014; Sundaram and Cohen, 2017).

How terminal cells form subcellular tubes can be studied live during embryonic development, when the apical membrane begins to invaginate while the first branch of the subcellular tube forms. The cytoskeleton is critical to drive this morphogenetic process at different steps. First, centrosomes localize subapically, from where they initiate lumen formation by organizing microtubules to pull the apical membrane inwards (Ricolo et al., 2016). Microtubules are also required for tube elongation; expression of Spastin, a microtubule-severing protein, prevents subcellular tube elongation (Gervais and Casanova, 2010). The actin cytoskeleton also plays an important role in tube formation. Conditions that disturb F-actin organization at the apical cortex result in shorter or misguided subcellular tubes, where the subcellular tube does not elongate in the direction of cell growth but does so in the backward direction (JayaNandanan et al., 2014). Crosstalk between microtubules and the actin cytoskeleton is also required for proper tube formation. Absence of Shortstop (Shot), a fly homolog of the F-actin-microtubule crosslinker Spectraplakin, also results in shorter subcellular tubes. Shot is thought to anchor the microtubules that surround the subcellular tube to an actin core that lies ahead of the subcellular tube, but whose function has remained elusive (Oshima et al., 2006; Ricolo and Araujo, 2020).

Tube formation and guidance are also affected by other actin-related processes that do not seem to be directly related to tube formation. Loss of the actin crosslinkers Fascin (encoded by *singed, sn*) or Forked (Espin) affects filopodia and terminal cell migration, but possibly as a secondary effect it also results in shorter or misguided tubes (Okenve-Ramos and Llimargas, 2014). Similar phenotypes are seen in cells lacking Enabled (Ena), a VASP-family actin polymerase involved in filopodia formation or Rhea, a Talin homolog that anchors integrins to the actin cytoskeleton (Levi et al., 2006; Gervais and Casanova, 2010; Choi et al., 2011). How and why disruption of the basal cortical actin meshwork in the filopodia is communicated to and influences the morphogenesis of the apical plasma membrane of the subcellular tube is an open question. Here, we address the role of actin organization at the tip of the cell, ahead of the tube, and propose a model where this compartment orchestrates distinct actin pools in the cell resulting in proper subcellular tube elongation and guidance. Our results point to a role for late endosomes as centres for actin nucleation ahead of the tube, a process that allows the crosstalk and coordination between distinct cytoskeletal pools within the cell.

## Results

### The late endocytic pathway and subcellular tube guidance

Late endosomes in tracheal cells act as a hub in the coordination of plasma membrane redistribution (Dong et al., 2014, 2013; Mathew et al., 2020). They are strongly enriched in the growth cone of tracheal terminal cells in the area between the extending filopodia and the tip of the growing subcellular tube (Mathew et al., 2020). These vesicles can be marked with CD4∷mIFP, a transmembrane construct that also labels the plasma membrane (Fig. 1A). They also carry Rab5 and Rab7, and we will refer to them collectively as late endosomes or CD4 vesicles. To study possible roles of their polarized distribution, we looked at loss-of-function conditions that affect late endosome formation. We showed previously that terminal cells mutant for *shrb* had large membrane aggregates as a result of failed recycling to the plasma membrane (Mathew et al., 2020). While a fraction of these cells was still able to form a tube, this was often abnormal, with the tube curling dorsally or forming an aberrantly branched tube (Fig. 1B-D, Movie S1). As an alternative way of perturbing endosomal maturation we expressed the dominant negative form of *Vha100*, *Vha100^R755A^*, a neural-specific subunit of the vesicular ATPase normally involved in endosomal maturation and acidification (Hiesinger et al., 2005; Williamson et al., 2010). Vha100^R755A^ blocks proton translocation into endosomes, therefore preventing their acidification. Its expression in larval terminal cells causes similar defects as loss of other vATPase subunits (Francis and Ghabrial, 2015). Expression of Vha100^R755A^ in embryonic terminal cells does not affect early endosome transition to late endosomes, as shown by the presence of FYVE∷GFP, a reporter for early endosomes and MVBs that localizes to CD4 vesicles both in control cells (Mathew et al., 2020) and in cells expressing Vha100^R755A^, which still formed large FYVE-positive vesicles towards the tip of the cell (Fig. S1A-B). But similar to loss of *shrb,* the expression of *Vha100^R755A^* in embryonic terminal cells also led to a higher proportion of cells with misguided tubes (Fig. 1C-D). This indicates that late endosome maturation is involved in proper subcellular tube guidance.

**Figure 1.**
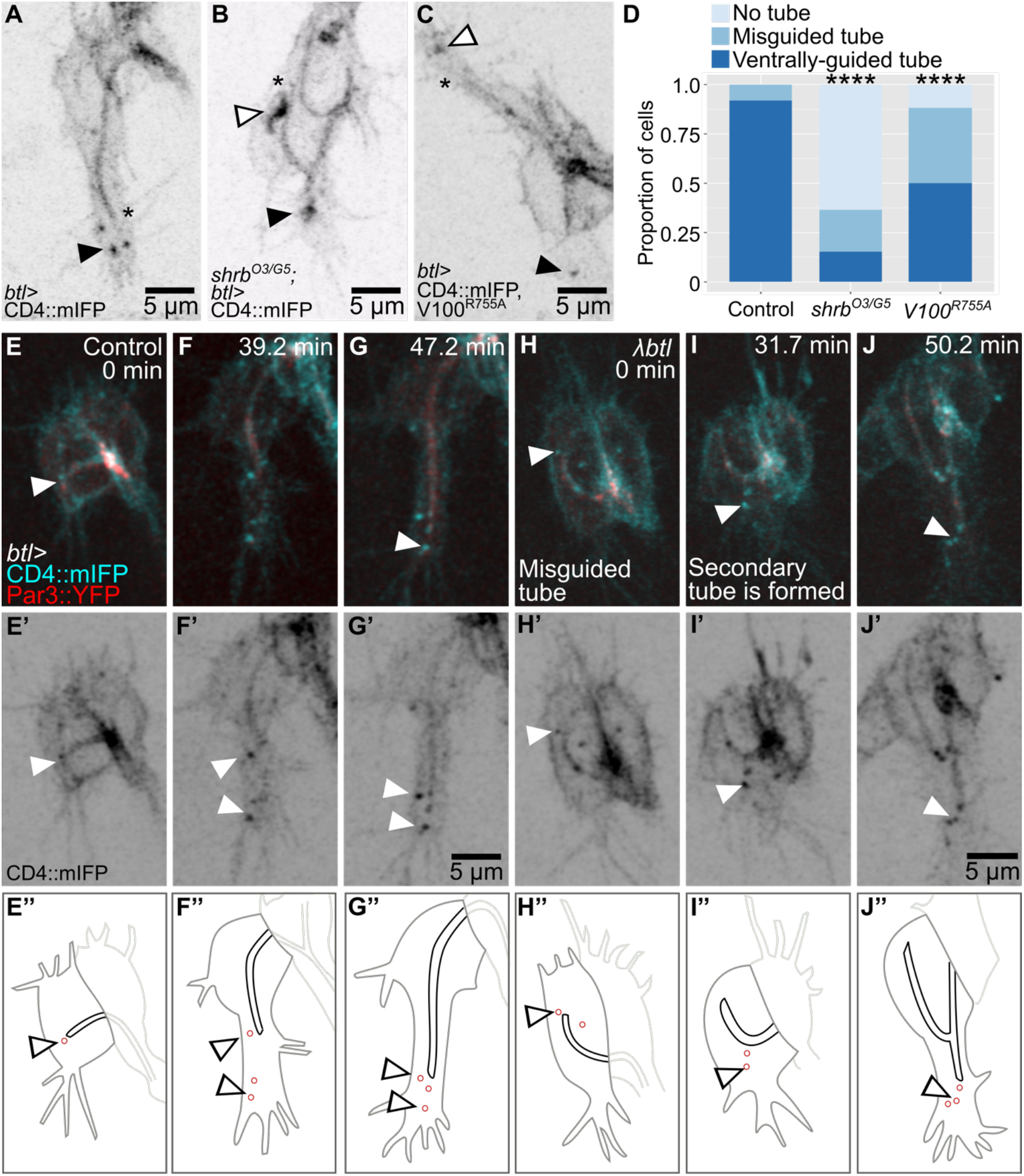
Distribution and role of late endosomes in subcellular tube guidance. (A-D) Terminal cells expressing CD4∷mIFP under *btl-gal4*. (A) Control. (B) Cell mutant for *shrb*. (C) Cell expressing *Vha100^R755A^,* a dominant negative construct that blocks vATPase function. (D) Proportion of cells with the indicated phenotypes. Number of cells analyzed: control, n=37; *shrb^O3/G5^*, n=33; Vha100^R755A^, n=42. Significance was assayed with a c^2^ test; ****p<0.0001. (E-J) Terminal cells expressing CD4∷mIFP and Par3∷YFP under *btl-gal4*. (E’-J’) CD4∷mIFP, (E’’-J’’) Manual tracings of the contours of the terminal cells (dark gray), subcellular tube (black), endosomes (red) and the adjacent dorsal branch cells (light gray). (E-G) Control. (H-J) Cells expressing a constitutively active FGFR, *λbtl*. Arrowheads point to CD4 vesicles.

To study the dynamics and location of late endosomes, we followed them in cells in which normal growth and branching were modified. In control cells, these vesicles can be seen as soon as the tube begins to form (Fig. 1E). As the cell and its subcellular tube elongate, CD4 vesicles remain associated with the growing tip of the cell (Fig. 1F-G). Excessive FGF signalling induces terminal cell duplication and misguided cell extensions (Lebreton and Casanova, 2016; Lee et al., 1996). Expression of *λbtl*, a constitutively active form of the FGF receptor, resulted in a range of defects that correlated with aberrant distribution of CD4 vesicles (Fig. 1IH-J, S1C-E). A cell with a curled-up tube also formed ectopic lamellipodia-like protrusions, and each of these contained CD4 vesicles (Fig. S1C). Duplicated terminal cells with branches that sprouted in aberrant directions also had CD4 vesicles in the space between the subcellular tube and the misguided growing tip of the cell (Fig. S1D).

Occasionally, terminal cells formed a secondary subcellular tube, and in these cases, the sprouting of the tube was preceded by the appearance of CD4 vesicles ahead of it (Fig. 1H-J, another example in S1E). The correlation of the presence of CD4 vesicles and ectopic branch and tube formation would also be consistent with a role of the vesicles in branch or tube morphogenesis or guidance.

### Mislocalization of late endosomes and tube guidance

It is also possible that the localization of these organelles at the tip of the cell is a secondary effect of signalling events but has no guiding function for tube growth. To test this we redirected vesicles carrying YFP-labelled Rab7 (YRab7), which in terminal cells is found almost exclusively in CD4 vesicles [Fig. 2A-C, (Mathew et al., 2020)] with GrabFP nanobodies (Harmansa et al., 2017). Specifically, we used GrabFP-B_Int_, a construct consisting of the anti-GFP nanobody vhhGFP4 which is marked with mCherry and fused to the transmembrane protein Nervana1 (Nrv1) with vhhGFP4 facing the cell cytoplasm (Fig. 2C). In imaginal discs, Nrv1 allows this construct to localize exclusively to the basolateral domain (Harmansa et al., 2017). However, in terminal cells, most of the protein failed to reach the basal membrane and instead formed membrane-associated aggregates within the cell (Figure 2A, C, Movie S2). Nevertheless, the construct served our purpose. We found that expressing GrabFP-B_Int_ in a YRab7 homozygous background resulted in association of GrabFP-B_Int_ and YRab7 puncta and it also increased the proportion of misguided cells compared to controls (Fig. 2E, G). In all cases where tube guidance was unaffected, cells retained a CD4 vesicle ahead of the tube, and this vesicle carried both GrabFP-B_Int_ and YRab7 (Fig. 2D).

**Figure 2.**
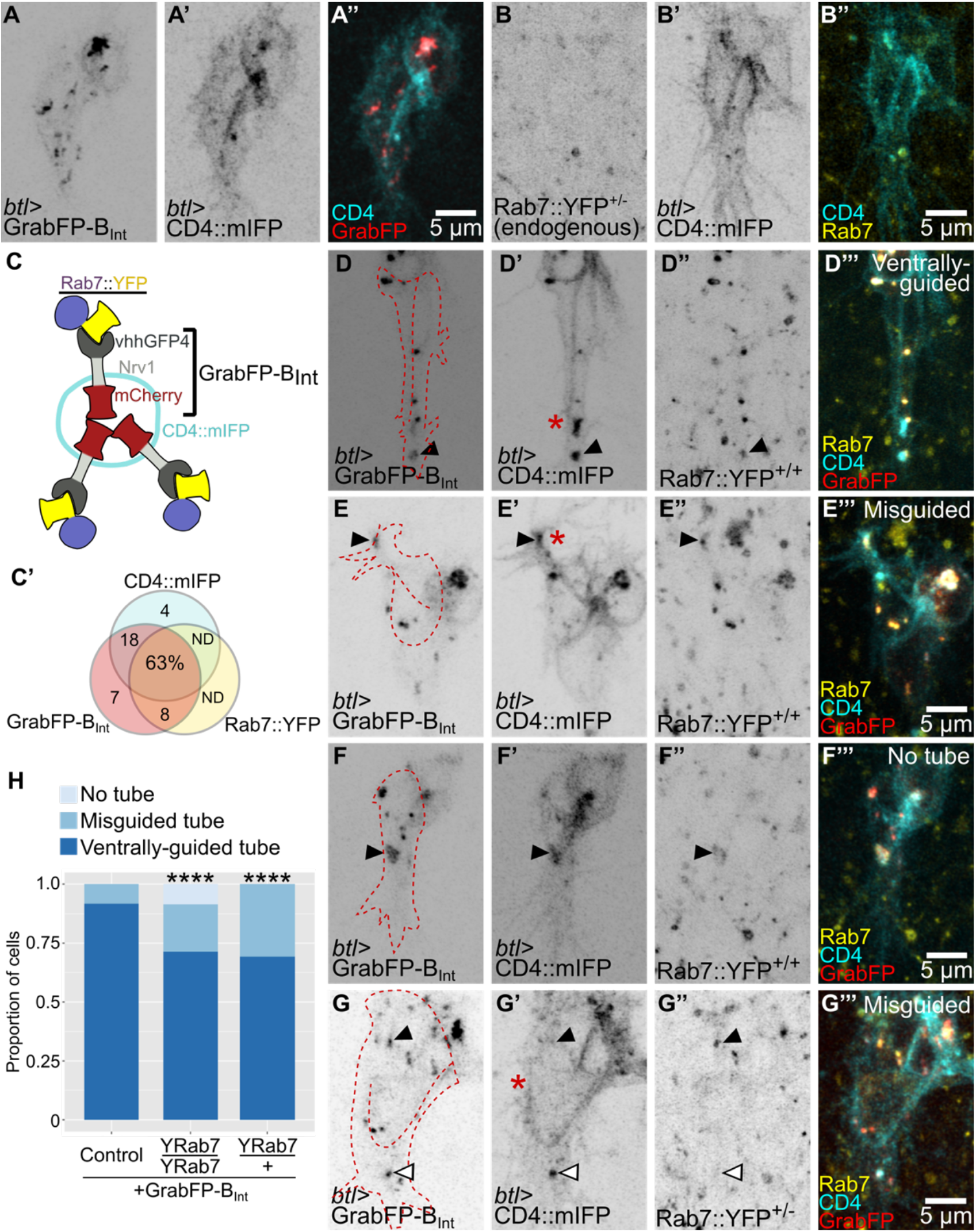
Mislocalization of late endosomes and subcellular tube guidance. (A-B) Terminal cells expressing CD4∷mIFP under *btl-gal4* together with GrabFP-B_Int_ (A) and endogenously labelled Rab7∷YFP (B). (C) Diagram of the GrabFP-B_Int_ nanobody in vesicles and associating with Rab7∷YFP around the vesicle. (C’) Venn diagram of vesicles containing CD4∷mIFP, GrabFP-B_Int_, or Rab7∷YFP. n=395 vesicles, from 6 different cells homozygous for Rab7∷YFP and collected across 7 time point each. ND = Not detected. (D-H) Cells expressing CD4∷mIFP and GrabFP-B_Int_ under *btl-gal4* in a Rab7∷YFP homozygous (D-F) and heterozygous background (G). The outline of the cells was traced using the CD4∷mIFP signal and is shown as a red dashed line. Black arrowheads point to vesicles containing CD4∷mIFP, GrabFP-B_Int_, and Rab7∷YFP, and white arrowheads point to vesicles that do not contain Rab7∷YFP. Asterisks mark the tips of the subcellular tubes in D’, E’, and G’. (H) Proportion of cells with the indicated genotypes. Number of cells analyzed: control, n=28; heterozygous YRab7 background, n=26; homozygous YRab7 background, n=59. Significance was assayed with c^2^ tests; ****p<0.0001.

Some cells had a particularly strong phenotype, where all detectable GrabFP-B_Int_, mCD4∷IFP and YRab7 formed a single aggregate and no tube was formed at all (Fig. 2F, H, Movie S2). This suggested that aggregation of YRab7 via GrabFP could block late endosome formation rather than affecting their distribution. However, expressing the construct in a YRab7 heterozygous background (where only one allele is sensitive to GrabFP) resulted in CD4 vesicles present at the tip of the cell but with no YRab7, and ectopic vesicles that carried GrabFP with the YRab7 protein (Fig. 2G, H). Despite the presence of presumably functional endosomes at the tip, an increased proportion of terminal cells had misguided tubes compared to controls. We conclude from these results that the correct distribution of late endosomes is required for proper tube guidance.

Further underlining the involvement of late endosomes in tube morphogenesis, hyperactivation of Rab7 was sufficient to affect branch and tube growth. Expression of Rab7^Q67L^, a GTP-locked, constitutively active form of Rab7, increases late endosome abundance in motor neurons and photoreceptors (Cherry et al., 2013), and we find that YFP-marked Rab7^Q67L^ (Zhang et al., 2007), while not detectable in the embryo when expressed under *btl-gal4*, led to significantly enhanced terminal cell branching in 3^rd^ instar larvae (Fig. S2A-B, D). Expression of a dominant negative form of Rab7 showed no obvious effect on larval branching (Fig. S2C, D), suggesting that terminal cells may not rely on late endosomes to build subcellular tubes at the larval stage, or that larval branching uses other compensatory mechanisms.

### Late endosomes and actin recruitment

The results so far suggested an instructive role of late endosomes in tube guidance, but it was not clear how this relates mechanistically to known regulators of this process, in particular the important function of the actin cytoskeleton (Oshima et al., 2006; JayaNandanan et al., 2014; Schottenfeld-Roames et al., 2014; Gervais and Casanova, 2010; Levi et al., 2006; Okenve-Ramos and Llimargas, 2014; Ricolo and Araujo, 2020). When we stained embryonic terminal cells expressing YRab7 with markers for actin (phalloidin-ATTO647) and the subcellular lumen (CBP-Alexa546), we found actin significantly enriched around YRab7 vesicles in the area ahead of the tube (Fig. 3A, B, S3). While actin was always seen enriched around these vesicles, its distribution varied from small puncta in the vicinity of the endosomes to thick networks that extended to the apical membrane surrounding the lumen (visualized with the subcellular lumen marker CBP; Fig. 3A, B, S3).

**Figure 3.**
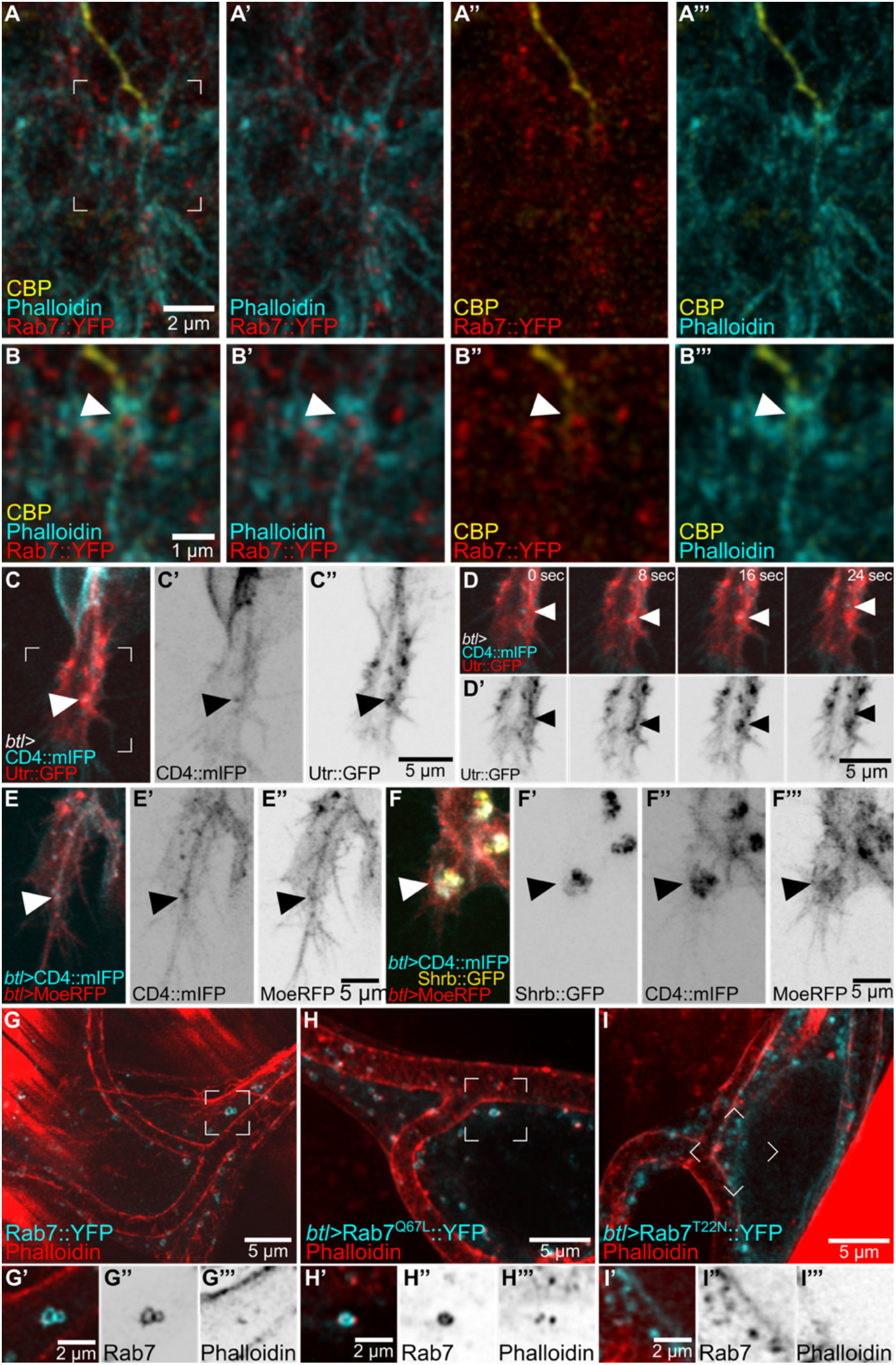
Actin organization at late endosomes. (A-B) Terminal cell expressing Rab7∷YFP and stained with phalloidin and a chitin binding protein (CBP) fused to Alexa647. (A-A’’’) Channels shown in pair-wise combinations. The region marked in A is magnified in B-B’’’. Arrowheads point to the contact between a Rab7∷YFP ring, actin and the tip of the subcellular tube. (C-D) Terminal cell expressing CD4∷mIFP and Utr∷GFP under *btl-gal4*. The square shown in C is magnified in D-D’ over four different time points. Arrowheads point to CD4 vesicles. (E-F) Terminal cells expressing CD4∷mIFP under *btl-gal4* and MoeRFP under direct control of the *btl* promoter. (E) Control, (F) cell also expressing Shrb∷GFP under *btl-gal4*. Arrowheads point to CD4 and MoeRFP puncta. (G-I) Larval terminal cells stained with phalloidin. (G) Control larva expressing endogenously labelled YRab7. (H) Cell expressing Rab7^Q67L^∷YFP. (I) Cell expressing Rab7^T22N^∷YFP. Regions marked in G-I are shown in G’-I’ but as single z-slices instead of projections.

We confirmed these results in live embryos expressing CD4∷mIFP and the actin-binding domain of Utrophin fused to GFP (Utr∷GFP), a faithful actin reporter with minimal effects on actin function (Spracklen et al., 2014). Utr∷GFP dynamically associated with CD4 vesicles near the tip of the cell (Fig. 3C-D, Movie S3). Another marker for actin, consisting of the actin-binding domain of Moesin fused to RFP (moeRFP), was seen in puncta in close proximity to CD4 vesicles (Fig. 3E). Furthermore, the large membrane aggregates with late endocytic markers caused by overexpressing Shrb∷GFP, as described above, had high levels of moeRFP enriched around them (Fig. 3F). We then turned to larval terminal cells, which have completed their branching morphogenesis but can serve as a system to study actin recruitment to late endosomes. We first looked at actin distribution relative to YRab7 using phalloidin in control larvae. Actin was often seen as small puncta associated with YRab7 vesicles (Fig. 3G). In terminal cells expressing the constitutively active Rab7^Q67L^∷YFP, actin puncta were associated with most of the Rab7^Q67L^∷YFP rings (Fig. 3H). In cells expressing a constitutively inactive form of Rab7, Rab7^T22N^∷YFP, actin did not associate with the labelled vesicles (Fig. 3I). In summary, late endosomes in terminal cells associate with and are able to recruit actin.

### Effect of tip laser ablation in terminal cell development

The observations so far suggest that actin may connect the apical and basal membranes at the tip of the cell to help the growing tube to follow the advancing growth cone. If this is the case then one might expect the actin to be under tension. To test this, we severed it by performing infrared laser cuts and measured recoil. We tested the ability of the laser to disrupt actin at a position where we could see the behaviour of the tube as a readout, approximately 1 micron behind the tip of the tube. This resulted in a rupture of the apical cortical actin cytoskeleton, with a herniation of the membrane as seen by CD4∷mIFP (Fig. 4A, Movie S4), which recovered after few seconds (Fig. 4B, Movie S4). Having confirmed an effect of the laser procedure on the cytoskeleton, we performed laser cuts ahead of the tube, and measured the initial recoil speed away from the cut using Particle Image Velocimetry (PIV) (Fig. 4C). Upon laser ablation, we observed a slight recoil in both directions (Fig. 4C, G, Movie S4). In normal control embryos, a small fraction of terminal cells produce a transient secondary branch that grows sideways or opposite to the direction of cell growth. This secondary branch usually also has late endosomes ahead, and therefore, also an additional actin core. We performed laser cuts in cells with these bifurcated tubes, and found that the initial recoil was faster than in cells with single, ventrally guided tubes (Fig. 4E, G, S4C-D, Movie S4). We also imaged longer-term cell recovery after laser manipulation. Cells with a single, properly guided tube were able to recover fast and resume elongation in the ventral direction (Fig. 4D, Movie S4). In contrast, in cells with tube bifurcations, the normal, ventrally growing tube initially recovered from the cut, but it subsequently collapsed (n=3/5 experiments; Fig. 4F, S4A-B, Movie S4). This response took place several minutes after the cut and cannot be explained by a direct effect of the laser cut. We propose this could be due to a collapse of the cytoskeleton in the affected tube. These experiments support the notion that adequate cytoskeletal organization at the tip is required for tube guidance and stability. The fact that cells with tube bifurcations have higher recoil speeds suggests that actin cores ahead of the tube generate tension to keep the tube in place, and that having more than one tip results in higher tension. Furthermore, the delayed collapse of bifurcated tubes suggests that the branches may be in some form of competition with each other.

**Figure 4.**
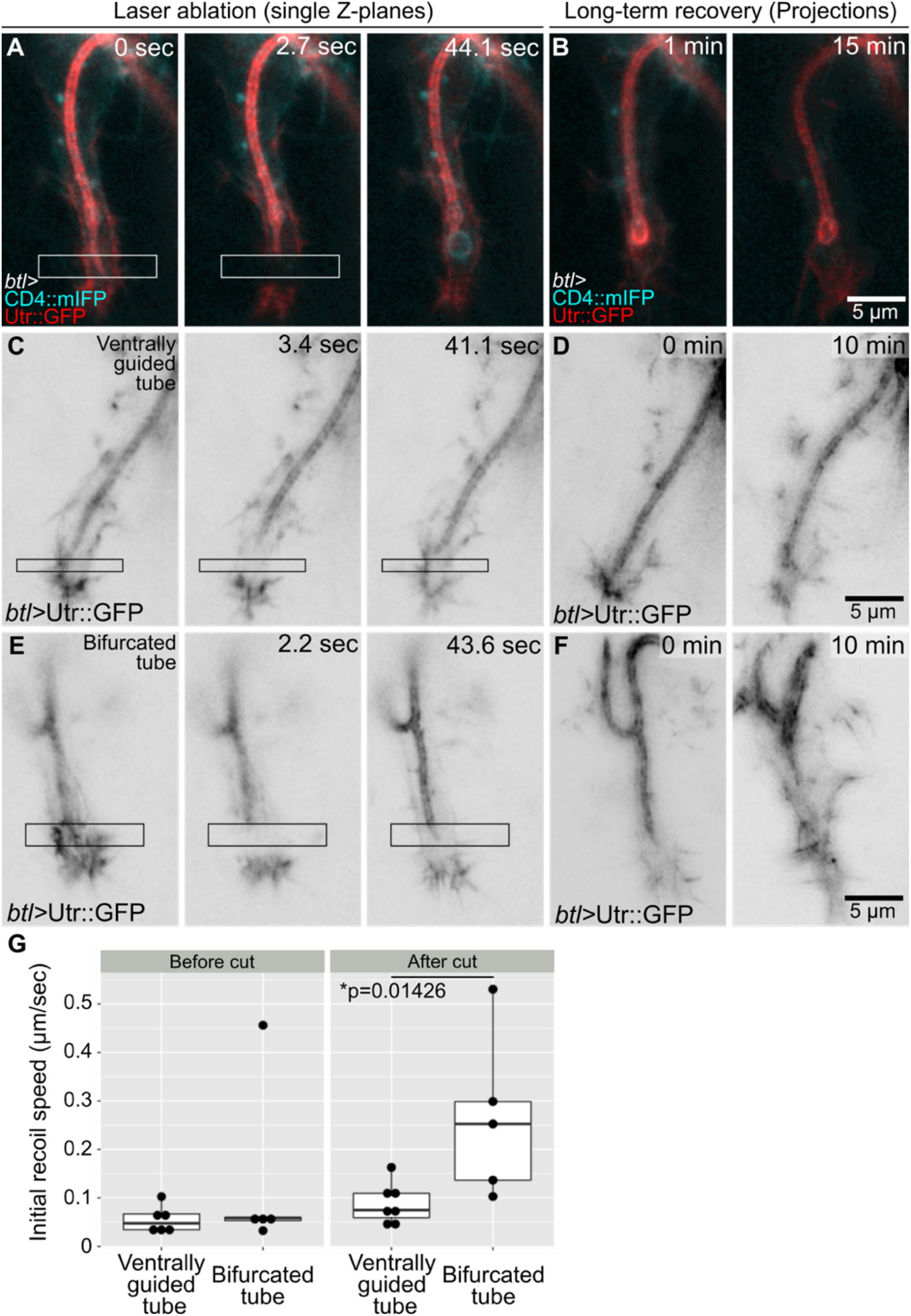
Infrared laser ablation at the growing tip of terminal cells. (A-B) Terminal cell expressing CD4∷mIFP and Utr∷GFP under *btl-gal4*. (A) Cell immediately before laser cut. The white box marks the laser-ablated region. (B) Long-term recovery, z-projections. (C-F) Terminal cells expressing Utr∷GFP under *btl-gal4.* (C-D) Cell with normal, ventrally guided tube, (E-F) Cell with a bifurcated tube. C and E are single z-plane images of cells immediately before laser cuts; black boxes mark the laser-ablated regions. D and F are z-projections of the recovery. (G) Quantification of the recoil as determined by particle image velocimetry (PIV). As control we analysed PIV in two successive timepoints before laser cuts. Number of cells analysed: cells with ventrally guided tube, n=7; cells with a bifurcated tube, n=5. Significance was assessed using t-test.

### The role of Wash in terminal cell development

Actin recruitment to late endosomes has been well documented in the context of recycling, where it is involved in budding of vesicles from endosomes, allowing retrieval of transmembrane cargo. This is mediated by Washout (Wash), a Wiskott-Aldrich Syndrome protein family member that activates the Arp2/3 complex thus allowing actin nucleation (Rottner et al., 2017; MacDonald et al., 2018; Gomez et al., 2012). Wash has been studied in multicellular tubes of the tracheal system, but its role in terminal cell development has not been determined (Dong et al., 2013; Tsarouhas et al., 2019). We found that a UAS-Wash∷GFP construct showed a strong, dynamic enrichment around CD4 vesicles (Fig. 5A). Knocking down *wash* in terminal cells carrying the PIP_2_-binding reporter PH∷GFP, to mark the subcellular tube membrane (Fig. 5B), resulted in tube misguidance phenotypes similar to those seen for disruption of endosomes (Fig. 5C, E). In addition, we noted that in cells where tube guidance was not affected the subcellular tube was abnormally short, with the distance between the tip of the tube to the tip of the cell significantly larger than in control cells (Fig. 5F). This phenotype is also seen upon the loss of the apical actin regulator Bitesize [Btsz, (JayaNandanan et al., 2014)].

**Figure 5.**
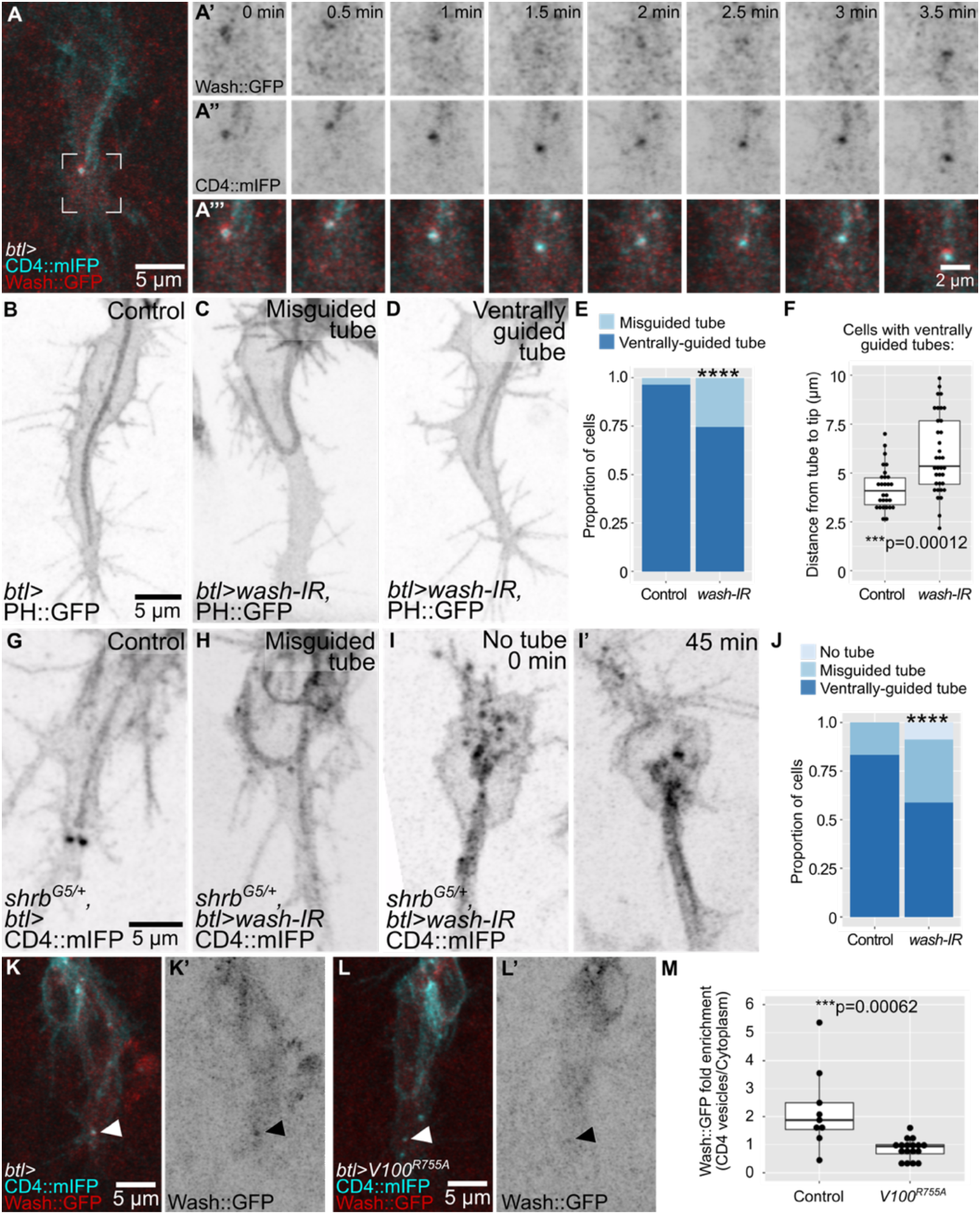
The role of Wash in terminal cell development. (A) Terminal cell expressing CD4∷mIFP and Wash∷GFP under *btl-gal4*. The region marked in A is shown in A’-A’’’ at higher magnification over eight time points. (B-F) Cells expressing PH∷GFP under *btl-gal4*; control (B) and *wash* knockdowns (C-D). (E) Proportion of cells with the indicated phenotypes. Number of cells analyzed: control, n=29; *wash-IR*, n=46. Significance was assessed with χ^2^ test, ****p>0.0001. (F) Distance between the tip of the tube to the tip of the cell in cells with a normal, ventrally guided tube. Number of cells analyzed: control, n=28; *wash-IR,* n=35. Significance was assessed with a Mann-Whitney U-test. (G-J) *shrb^G5^* heterozygous embryos expressing CD4∷mIFP under *btl-gal4*; *shrb^G5^* heterozygote as control (G) and *shrb^G5^* heterozygote expressing *wash-IR* (H-I). (H) cell with a misguided tube. (I) Cell lacking a tube. (J) Proportion of cells with the indicated phenotypes. Number of cells analyzed: *shrb^G5^* heterozygotes (controls), n=30; *shrb^G5^* heterozygotes expressing *wash-IR*, n=34. Significance was assessed with χ^2^ test, ****p>0.0001. (K-M) Cells expressing CD4∷mIFP and Wash∷GFP under *btl-gal4*. (K) Control, (L) cell also expressing Vha100^R755A^. Arrowheads point to CD4 vesicles. (M) Fluorescence intensity of Wash∷GFP at CD4 vesicles normalized over the mean signal of Wash∷GFP in the cytoplasm. Number of cells analyzed: control, n=11; Vha100^R755A^, n= 17. Significance was assessed with a Mann-Whitney U-test.

*wash* loss-of-function phenotypes could be due to a role not related to late endosomes, like actin nucleation in a different cell compartment. To test this, we analysed genetic interactions between *shrb* and *wash* by expressing *wash* RNAi in a *shrb* heterozygous background. We found that the proportion of cells with misguided tubes was higher upon *wash* knockdown compared to control *shrb* heterozygotes (Fig. 5G-J). Furthermore, in the *shrb*^+/−^, *wash-RNAi* condition we found a fraction of cells where no tube was formed at all (Fig. 5I-J). Wash is recruited to late endosomes upon acidification (Nagel et al., 2017), and since we had found that expression of Vha100^R755A^ (which prevents endosome acidification) leads to tube misguidance, we wondered if this was due to a failure to recruit Wash∷GFP to late endosomes. We found that in cells expressing Vha100^R755A^, Wash∷GFP recruitment was significantly less prominent than in control siblings (Fig. 5K-M). The number and positioning of the vesicles was not affected by the perturbations (Fig. S5F).

### Actin distribution upon loss of components of the late endocytic pathway

To test if actin recruitment to late endosomes depends on Wash, we quantified the fluorescence intensity of Utr∷GFP that overlapped with CD4 vesicles in cells expressing Vha100^R755A^ or *wash* RNAi, with similar results in both cases. Compared to controls (Fig 6A-B, Movie S5), CD4 vesicles had significantly less Utr∷GFP fluorescence intensity associated with them,. In both conditions, actin organization in other areas of the growth cone also seemed to be affected more broadly (Fig. 6C-F, S5E, Movie S5). Both the apical actin mesh surrounding the tip of the tube and the basal actin cortex appeared reduced in this area but not in other parts of the cell. Since expression of Vha100^R755A^ is unlikely to affect the actin directly, and since the effect is restricted to the area around the leading endosomes, this suggests that the reduction in cortical actin is a secondary effect of actin reorganisation around the endosomes. In addition, within each experimental condition, we found a correlation between Utr∷GFP fluorescence intensity at CD4 vesicles and tube guidance: cells with less Utr∷GFP associated with the vesicles were more likely to have misguided tubes than cells of the same genotype that had normal actin levels around the vesicles (Fig. 6G). These results support a role of late endosomes in organizing actin, a property that is impaired in the absence of Wash or proper endosome acidification. Based on this, and on the phalloidin stainings in control embryos and the laser ablations, we propose that late endosomes at the tip serve as actin-organizing centres that contribute to the formation of a continuous actin meshwork from the tube to the basal plasma membrane. This model explains why an intact actin pool at the basal plasma membrane is required for proper tube guidance.

**Figure 6.**
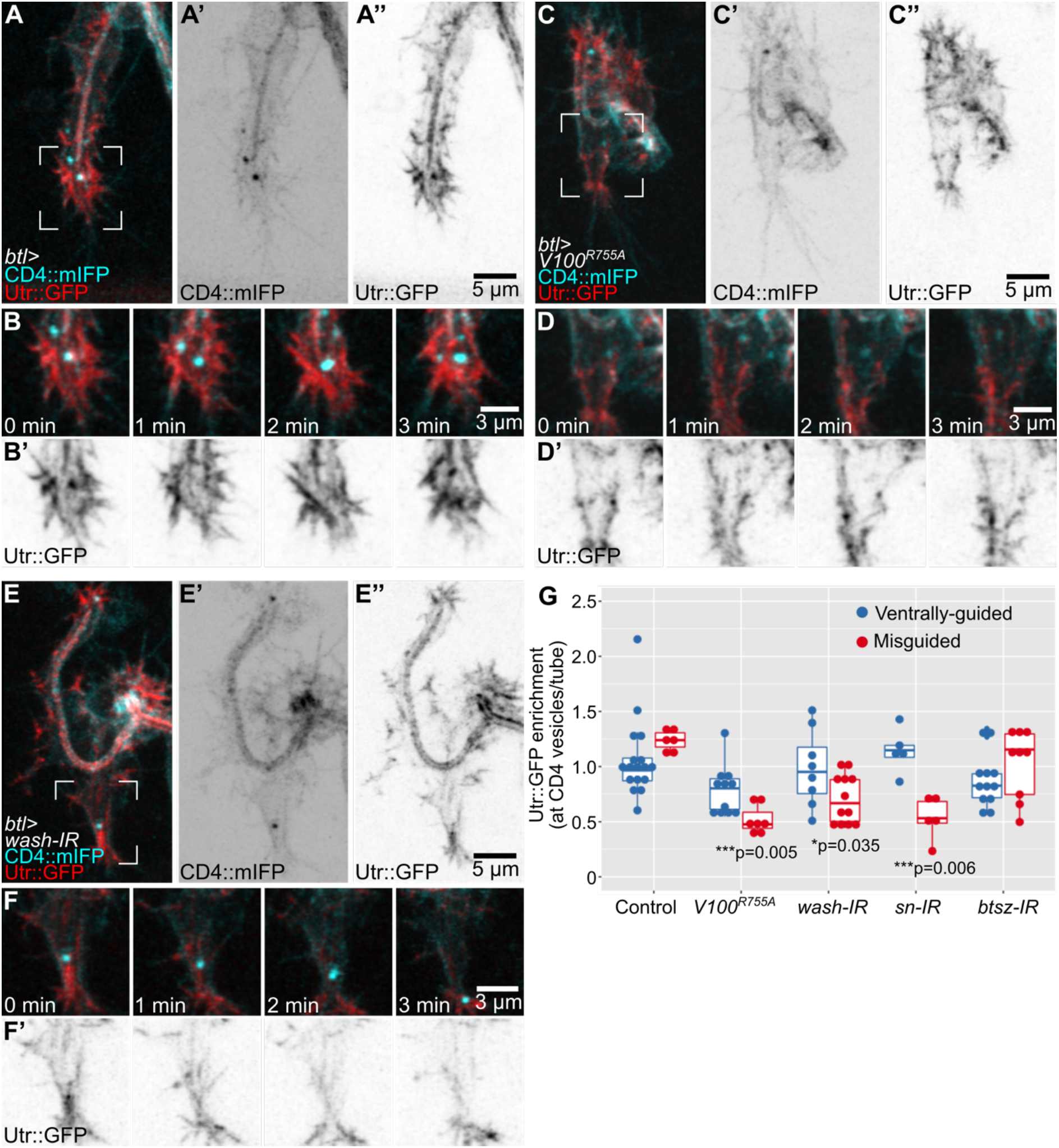
Actin organization in cells with defective late endosomes. (A-H) Cells expressing CD4∷mIFP and Utr∷GFP under *btl-gal4*. (A-B) Control, (C-D) cell expressing Vha100^R755A^, (E-F) *wash* knockdown. Regions marked in A, C and E are magnified and shown over four time points in B, D and F, respectively. (G) Mean Utr∷GFP fluorescence intensity in CD4 vesicles over time. Values were normalized to fluorescence intensity at the tube to adjust for variations in the expression of the Utr∷GFP reporter. Number of vesicles analyzed: control, n=23 from 9 cells; Vha100^R755A^, n=19 from 8 cells; *wash-IR*, n=20 from 9 cells; *sn-IR,* n=10 from 5 cells; *btsz-IR*, n=20 from 8 cells. Values at the bottom are significance between normal vs misguided cells within each genotype, assessed by t-test.

According to this model, the previously described guidance defects seen after the loss of actin regulators in the apical or basal cortices would be at least partly due to lack of continuity of the actin meshwork between the two compartments, with late endosomes acting as a ‘relay’ point. To test this, we analysed CD4 vesicles and their surrounding actin mesh in cells in which either the apical or basal meshwork had been disrupted by downregulating Btsz or the actin bundler Fascin, respectively (JayaNandanan et al., 2014; Okenve-Ramos and Llimargas, 2014). We found that these perturbations had different consequences. In cells expressing *sn* (Fascin) RNAi we found a significant reduction of Utr∷GFP at CD4 vesicles, and the absence of actin around vesicles correlated with the degree of misguidance of the tube, suggesting that apical misguidance may not be a direct consequence of basal actin loss, but was mediated via the CD4 vesicles (Fig. 6G, S5A-B, E). Conversely, in cells expressing *btsz* RNAi (Fig. 6G, S5C-E), Utr∷GFP was still associated with CD4 vesicles at similar levels as in controls. This was also the case for cells with misguided tubes (Fig. 6I). Thus, even though F-actin forms around CD4 vesicles, this meshwork cannot guide tube growth when the apical cortical actin at the tube is disrupted due to the absence of Btsz. Since the vesicles are still found at the growing tip of the cell, this experiment also suggests that their positioning is not dependent on the apical actin cytoskeleton.

## Discussion

Crosstalk between cortical actin pools is important for proper morphogenesis in several developmental contexts (Goodwin et al., 2016), and we describe a mechanism by which this can be achieved. Late endosomes at the growing tip of terminal cells act as platforms to assemble cytoskeletal regulators or signals that propagate forces within the cell. This dynamic actin meshwork likely connects other cytoskeletal components, like the more stable actin pools at the apical and basal cortices of the cell, thereby coupling the directionality of growth of the two membrane compartments. Loss of actin regulators at any of these structures uncouples the direction of tube growth from that of cell elongation, resulting in tube misguidance (Fig. 7).

**Figure 7.**
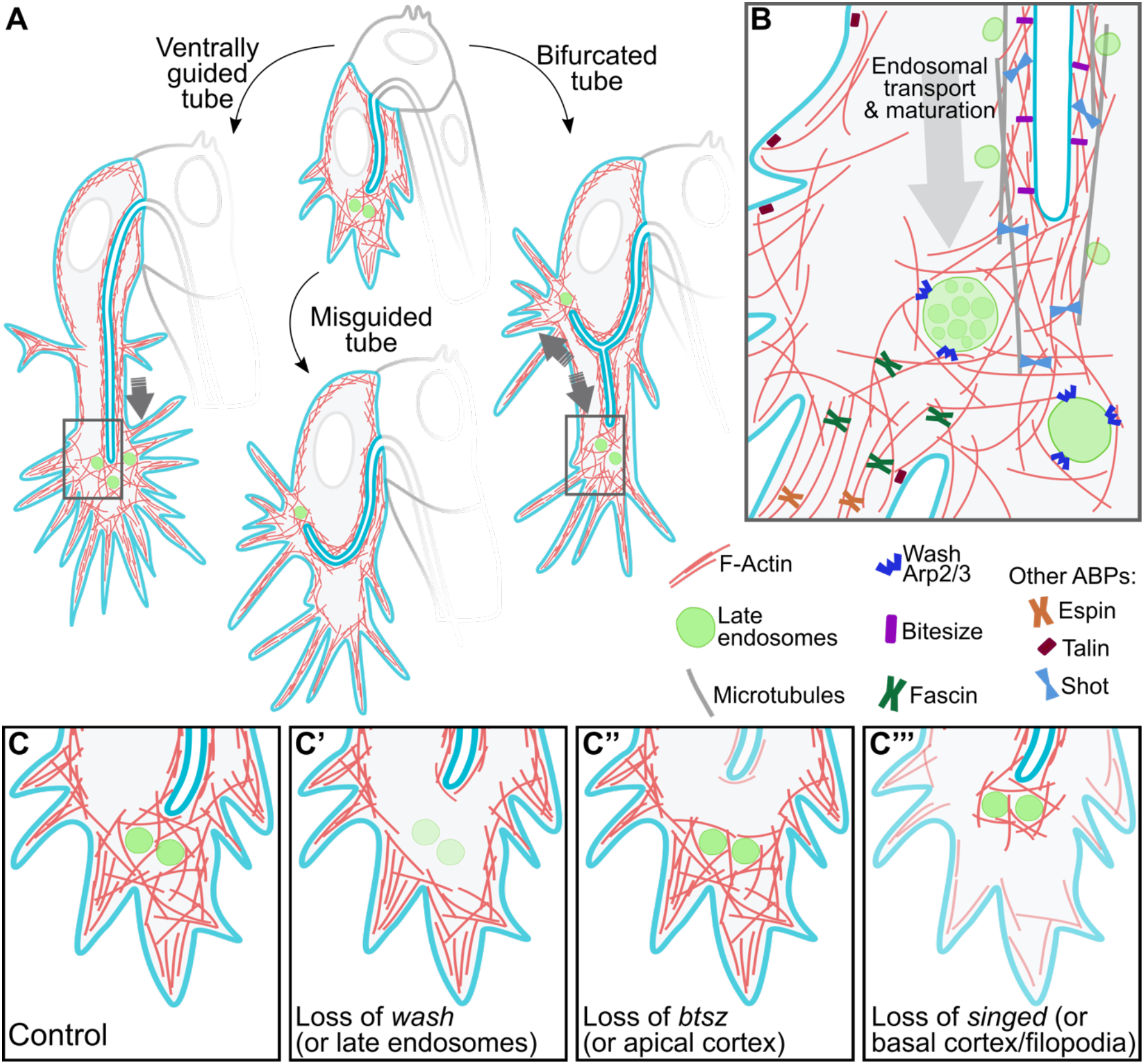
Graphic summary of the role of late endosomes in subcellular tube guidance. (A) Control cells. Late endosomes situated at the tip of the cell allow the organization of an actin network that extends from the tube to the growing tip of the cell. Control cells with misguided tubes result from the nucleation of actin surrounding late endosomes not located at the tip. In cells with tube bifurcations, late endosomes form two actin nucleation centers that stabilize each subcellular branch. (B) Zoom-in to the tip of the cell (regions marked with rectangles in A). Distinct actin-binding proteins (ABPs) at different subcellular compartments coordinate proper tube formation, with Wash acting in late endosomes to bridge actin pools at the apical (via Bitesize) and basal (via Talin and the filopodial organizers Fascin and Espin) cortices. Also shown are microtubules that surround the subcellular tube and that are anchored to the actin meshwork at the tip via Shot. (C-C’’’) Effects of loss of actin organizers at distinct subcellular compartments. (C) Control. (C’) Upon loss of *wash* or in conditions with aberrant endosomal maturation, F-actin fails to form around late endosomes. (C’’) In the absence of *btsz* or other actin organizers at the apical cortex, F-actin fails to crosslink to the actin meshwork at the tip. (C’’’) Lack of actin regulators at the basal cortex prevents the connection of the actin meshwork at the tip to the basal plasma membrane. All these conditions decouple the growth of the tube from that of the tip of the cell.

### The late endocytic pathway in tube formation

Our previous analyses of the late endocytic pathway in terminal cells led us to propose that endosomes derived from the subcellular tube undergo maturation as they travel to the tip of the cell, from where material is sorted towards the basal plasma membrane or recycled back to the apical domain (Mathew et al., 2020). This is a very dynamic process in which new vesicles constantly reach the tip of the cell and fuse with existing mature endosomes. Here we show that these endosomes also recruit Wash, and our current experiments and previous work suggest Wash recruitment depends on endosome acidification (Nagel et al., 2017). Wash function in late endosomes has been linked to protein recycling, in particular of integrins (Nagel et al., 2017; MacDonald et al., 2018). In tracheal multicellular tubes, Wash is involved in apical recycling (Dong et al., 2013). Our model agrees with these results, where late endosomes sort material destined to the apical and the basal plasma membranes. These results could argue for a failure in delivery of the FGF receptor to the basal plasma membrane as the cause of tube misguidance phenotypes. However, we also show here that increasing late endosome formation (by expression of Rab7^Q67L^) or mis-localizing late endosomes using GrabFP is sufficient to alter tube formation. This points to a specific role of late endosomes in tube guidance, rather than an indirect effect of defective FGF signalling.

Our results also show that while complete loss of the endocytic pathway prevents tube formation either in single mutants or in genetic interaction experiments, partial loss of a functional endocytic pathway leads to subcellular tube defects like misguidance, with little or no effect on early steps of endosome maturation. We assume that partial loss of function of the endocytic pathway still allows material to reach both plasma membrane compartments but it prevents proper actin organization around late endosomes, uncoupling the growth direction of the tube and the cell. Others have shown that loss of components of the vATPase, as well as loss of ESCRT-0 components, affect endosome and subcellular tube formation in larval terminal cells (Francis and Ghabrial, 2015; Chanut-Delalande et al., 2010; Nikolova and Metzstein, 2015). We interpret these results at least partially as a consequence of cytoskeletal disorganization near or at late endosomes, that leads to failure in tube formation.

Actin nucleation at endosomes has been identified as a mechanism that allows proper cell shape changes in other systems. In response to hepatocyte growth factor (HGF), cultured hepatocytes form endosomes that recruit and activate Rac. Activated Rac is then transported while on endosomes to the leading edge of the cell, where it favours the formation of protrusions that allow cell migration (Palamidessi et al., 2008). Also, lysosomal transport to the cell periphery has been implicated in focal adhesion dynamics, where peripheral transport of lysosomes allows the turnover of specific focal adhesion components (Schiefermeier et al., 2014). These studies highlight multiple roles of late endosomes in cell migration and actin organization.

### Positioning of late endosomes to the growing tip of terminal cells

What cues lead to the tip localization of late endosomes is still an open question. Late endosomes are transported along microtubules, so one possibility is that the direction of microtubule growth is defined by signalling mechanisms like the FGF signalling pathway, which also dictates the direction of cell growth. An alternative model is that microtubule organization and therefore transport of late endosomes is an indirect consequence of cell growth. Gervais and Casanova (2010) have proposed that the actin cytoskeleton, downstream of FGF signalling, allows polarized microtubule organization, and the distribution of Shot, an actin-microtubule crosslinker, supports this model (Ricolo and Araujo, 2020). Several studies have shown that multiple cytoskeletal regulators lie downstream of FGF signalling activation, namely, Ena (VASP family founding member), Fascin/Singed, and Shot itself (Gervais and Casanova, 2010; Okenve-Ramos and Llimargas, 2014; Ricolo and Araujo, 2020). The actin cytoskeleton is widely used to control guidance of microtubules (Dogterom and Koenderink, 2019), so it is reasonable to assume that FGF signalling could control microtubule orientation by locally organizing F-actin at the leading edge, which in turn serves as a cue to organize microtubules via Shot.

In human cells as well as in fly fat body cells and neurons, MVBs and lysosomes are constitutively transported to the cell periphery. This process uses microtubules and the ADP ribosylation factor (Arf)-like family member Arl8 (Rosa-Ferreira et al., 2018; Boda et al., 2019). It is possible that terminal cells use a similar mechanism to localize late endosomes to the growing tip of the cell, adopting a conserved, widespread process of peripheral positioning of late endosomes for a cell morphogenetic mechanism. Given that late endosomes are also required to deliver membrane and proteins to the basal membrane compartment (Mathew et al., 2020), peripheral late endosomal transport would also provide material to the growing tip of the cell.

### Crosstalk between cytoskeletal compartments

For late endosomes to propagate morphogenetic forces within the cell, they should be able to form an actin meshwork that is connected to other subcellular compartments. Our results show that the location of late endosomes correlates with the local formation of lamellipodium-like protrusions in the basal plasma membrane, for instance, in experiments where we over-activate FGF signaling. Similarly, affecting endosomal maturation has a general effect on actin organization throughout the tip of the cell. Interaction between actin pools has also been observed in multicellular tracheal tubes, where Wash activity balances the organization of endosomal and cortical actin pools (Tsarouhas et al., 2019). While Wash recruitment can explain the nucleation of actin at late endosomes, it does not explain how this network is crosslinked to the apical and basal actin cortices. Among the known actin regulators expressed in terminal cells, IKK*ε* and Singed/Fascin are good candidates to carry out such a function (Oshima et al., 2006; Okenve-Ramos and Llimargas, 2014). Both molecules are present throughout the cell cytoplasm, unlike others that show specific subcellular localizations, like Ena (Gervais and Casanova, 2010). Furthermore, loss of either IKKe or Singed/Fascin leads to general actin disorganization (Oshima et al., 2006; Okenve-Ramos and Llimargas, 2014), and our results using *sn* RNAi show that actin decrease at the tip correlates with subcellular tube misguidance. While myosin is dispensable for tracheal morphogenesis (Ochoa-Espinosa et al., 2017), actin crosslinking alone is sufficient to generate tension in other contexts (Bun et al., 2018), so it is likely that the action of these molecules connects the actin cortices with the endosomal actin and drives tube guidance in response to signalling.

Another key component involved in tube formation are microtubules, which initiate tube membrane invagination and allow further tube branching (Ricolo et al., 2016; Ricolo and Araujo, 2020). Microtubules are stiffer and less dynamic than actin, so it is likely that their role in tube formation is the stabilization of the membrane invagination as it continues to grow. In terminal cells, microtubules line the subcellular tube, with newer microtubules localized towards the growing tip (Gervais and Casanova, 2010). Increasing the number of microtubule-organizing centres (MTOCs) leads to the formation of ectopic subcellular tubes, and ahead of each tube, an actin core is also formed (Ricolo et al., 2016). Conversely, loss of microtubule regulators leads to shorter or lack of subcellular tubes, but not to tube misguidance (Gervais and Casanova, 2010; Ricolo et al., 2016; Ricolo and Araujo, 2020). These observations support the model where microtubules provide a mechanical structure that stabilizes tube growth, and at the same time they allow the transport of late endosomes to the tip of the cell. Late endosomes at the tip reinforce microtubule anchoring to the actin cytoskeleton, thereby linking tube stabilization and guidance in response to signalling cues. Further support for this model is provided by the fact that Wash can interact with tubulin in mammalian cell culture experiments (Derivery et al., 2009). Direct interaction between Wash and microtubules could be an additional factor that allows bridging the subcellular tube to late endosomes ahead of it. This would explain why expression of Rab7^T22N^ in larval terminal cells does not affect branching even though it prevents actin recruitment at late endosomes; microtubules might still be able to bind to late endosomes as a compensatory mechanism via Wash.

If microtubules play a more stabilizing role during tube growth, our observations of tube collapse several minutes after laser ablation could be explained by a failure in microtubule stability, caused by ablation of the actin meshwork that should normally anchor microtubules to the tip. In other systems, like the leading edge of migrating cells, neuronal growth cones and formation of dendritic spines, actin filaments open the way for the formation of new protrusions; these protrusions are then mechanically stabilized by microtubules (Dogterom and Koenderink, 2019). In the subcellular tubes of the nematode excretory system, microtubules also line the subcellular tube and they are anchored at the growing tip via formins to the actin cytoskeleton, and MVBs have also been found in this region of the cell (Sundaram and Cohen, 2017; Shaye and Greenwald, 2015). Coupling of cytoskeletal components by late endosomes as presented in this work could represent a widespread mechanism for cell morphogenesis.

## Methods

### Fly stocks

All UAS constructs were expressed using *btl-gal4* on the 2nd or 3rd chromosome (Chanut-Delalande et al., 2007; Shiga et al., 1996). UAS*-Utr∷GFP* and *UAS-PLCδ-PH∷GFP* are from Thomas Lecuit, IBDM, France, and we have used them for tracheal cells previously (JayaNandanan et al., 2014; Mathew et al., 2020). *UAS-PLCδ-PH-Cherry* is located at the VK00033 site and was cloned using *UAS-PLCδ-PH∷GFP* as a template (Mathew et al., 2020). The following stocks are from Bloomington Drosophila Stock Center: *UAS-Rab7^T22N^∷YFP* (#9778), *UAS-Rab7^Q67L^∷YFP* (#9779), *UAS-shrbGFP* (#32559), *shrb^O3^* (#39623), *shrb^G5^* (#39635), *UAS-GFP-myc-2xFYVE* (#42716), *UAS-CD4∷mIFP-T2A-HO1* (#64182, #64183), and *UAS-GrabFP-BInt∷mCherry* (#68175); and *UAS-sn-IR* (#32579), *UAS-btsz-IR* (#35205) and *UAS-wash-IR* (#24642) are from the Vienna Drosophila Resource Center.

We are grateful to the groups that kindly shared the following lines: UAS-Wash∷GFP from Sven Bogdan, Philipps-University Marburg, Germany (Nagel et al., 2017); endogenously YFP-labeled Rab7 (*YRab7*), from Marko Brankatschk, TU-Dresden, Germany (Dunst et al., 2015); *UAS-3xeYFP-Baz* from Chris Doe, University of Oregon, USA (Siller et al., 2006); *UAS-Vha100^R755A^* from Robin Hiesinger, Free University Berlin, Germany (Williamson et al., 2010) and *UAS-λbtl* from Denise Montell, UCSB, USA (Lee et al., 1996).

### Staining of embryos

Embryos were dechorionated as above and then fixed using 10% methanol-free formaldehyde and heptane for 20 minutes. Afterwards embryos were hand-devitellinized and blocked in a solution containing PBS, Triton X-100 0.5% and BSA 1% (Rothwell and Sullivan, 2007). Embryos were then stained with a solution containing 1:100 chitin-binding protein fused to Alexa564 (JayaNandanan et al., 2014), 1:200 phalloidin conjugated to ATTO647 (gift from Wioleta Dudka, EMBL), and 1:300 FITC-conjugated goat GFP antibody (Abcam #ab6662). Embryos were then washed and mounted in Vectashield. Images were acquired in a Zeiss LSM 880 in confocal mode using a Plan-Apochromat 63x/1.4 Oil objective, deconvolved in Huygens Professional and processed in FIJI.

### Staining of larvae

For branch number analyses, third instar larvae were heat-fixed by incubating them at 65°C for 15 seconds in halocarbon oil. Animals were then mounted with the dorsal side facing down and imaged in a Zeiss LSM 780 with a Plan-Apochromat 20x/0.8 M27 objective. Branches were manually counted from Z stacks in FIJI.

Larvae to be stained were dissected in PBS while immobilizing them with pins, cutting through the ventral midline. Animals were fixed using PFA 4% for 20 minutes, internal organs removed, and the resulting filets were washed with PBS Triton X-100 0.3% and blocked in PBS, Triton X-100 0.3%, BSA 1%. Animals were then stained with phalloidin conjugated to ATTO647 and FITC-conjugated GFP antibody (as above). We acquired images using a Zeiss LSM 880 in super-resolution mode and a Plan-Apochromat 63x/1.4 Oil objective. These were deconvolved using Zeiss Zen with automatic settings, and further processed in FIJI.

### Live imaging

We collected embryos in agar plates. Embryos were then dechorionated with 50% bleach, washed, and mounted with Halocarbon oil in glass-bottom dishes covered with heptane glue. Imaging was done in a Zeiss LSM 880 in Airyscan fast mode using a Plan-Apochromat 63x/1.4 Oil objective. Images were deconvolved in the Zeiss Zen software using auto settings, and further processed in FIJI. We prepared the movies using FIJI and iMovie.

### Laser ablation

Embryos were processed for live imaging as above. Ablation experiments were carried out in a Zeiss LSM 780 microscope equipped with a femtosecond two-photon laser (Chameleon Compact OPO Family, Coherent) at 950nm. To define the region of ablation, we used the Zen Bleaching module. Ablation was done as one cycle with 75% laser power (1540 mW total power) in single-z plane settings, whereas for the recovery we imaged z-stacks.

### Image analyses

#### Pixel intensity correlation

We used the Coloc2 plugin from FIJI to determine the degree of overlap between phalloidin and YRab7 signals using the Pearson’s correlation coefficient. We compared the results obtained from maximum intensity projection images of complete terminal cells with single-z close-ups to the tip of the cell where late endosomes are present.

### Branch counts and vesicle composition

We used FIJI to manually count branches from larval terminal cells from z-stacks. To determine the composition of vesicles in the nanobody experiments, we manually added the vesicles seen in each channel to the ROI manager and used this to compare whether a given vesicle also contained any other marker.

### Tube to tip of the cell measurements

We used FIJI to manually measure the distance between the tip of the tube to the growing tip of the cell. We used the line function from the tip of the tube to the farthest position of the tip of the cell, at the base of filopodia. All measurements were done at cells after ~1 hour after the tube began to grow.

### Wash and actin recruitment at CD4 vesicles

Using FIJI, we did SUM z-projections, and then manually subtracted the background from regions outside of the cell in the Wash∷GFP or Utr∷GFP channels, respectively. We manually segmented CD4 vesicles using the circle function, selecting vesicles that could be traced towards or at the tip of the cell for at least 3 timepoints and up to ~15. Values obtained from a single CD4 vesicle were then averaged. To normalize to the total amount of the reporters in the cell, we manually segmented the tube (for Utr∷GFP) or the cytoplasm (for Wash∷GFP) in three different timepoints where the corresponding vesicle was also present.

### Recoil measurements

We used the Particle Image Velocimetry (PIV) plug-in in FIJI to estimate recoil speeds (Tseng et al., 2012). We defined two 70×60 px regions of interest, one ahead and one behind the ablated area, which we defined as ‘forward’ and ‘backward’ recoil areas. Our PIV parameters were 20px for interrogation window size and 40px for search window size. We took the magnitude results from both regions of interest and then averaged the absolute values. As controls, we used the same regions but one time point before the laser cut. In addition to this analysis, we also measured recoil speeds in FIJI using the manual tracking plug-in. For this, we used three measuring points: the backward displacement of (i) the subcellular tube and (ii) of the plasma membrane adjacent to the ablated area, which both retract towards the cell body, and we also measured (iii) the forward displacement of plasma membrane on the other side of the ablated area. We then calculated the average of these for each experiment.

### Statistics

We used GraphPad Prism and QuickCalcs for statistical analyses. Plots were generated in RStudio using ggplot2.

## Acknowledgements

We thank Sofia Araujo, Phillipe Bun, Benedikt Best and Renjith Mathew for insightful comments on the manuscript; Sourabh Bhide for assistance on the laser ablation experiments, Juan Gomez for assistance in manual embryo devitellinization, and the entire Leptin lab for critical discussions. We thank Sven Bogdan for suggesting the Wash experiments. We are grateful to Robin Hiesinger, Sven Bogdan, BDSC and VDRC for providing fly stocks. We thank the EMBL Advanced Light Microscopy Facility (ALMF) for their support and Zeiss for the support of ALMF. This work was supported by funding from EMBL and EMBO. LDRB was funded by the EMBL Interdisciplinary Postdoctoral Programme under Marie Curie Actions.

## Author Contributions

Conceptualization, L.D.R.B.; Methodology and investigation, L.D.R.B.; Formal Analysis, L.D.R.B.; Writing – Original Draft, L.D.R.B.; Writing – Review & Editing, L.D.R.B., M.L.; Visualization, L.D.R.B.; Supervision, M.L.; Funding Acquisition, M.L.

---

**Figure S1.**
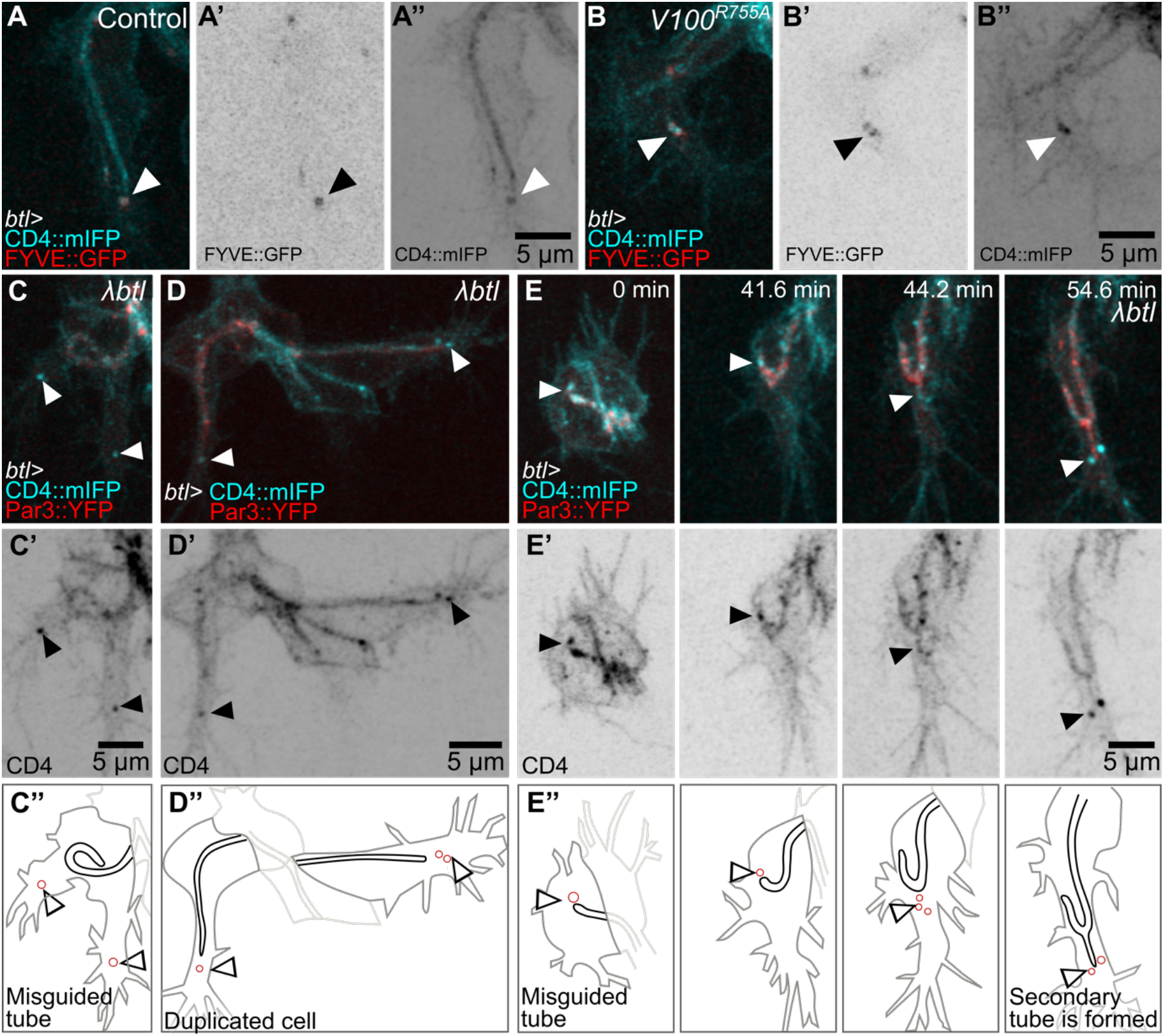
Endosome maturation and distribution during subcellular tube growth. (A-B) Terminal cells expressing CD4∷mIFP and FYVE∷GFP under *btl-gal4*. (A) Control. (B) Cells additionally expressing Vha100^R755A^. (C-E) Terminal cells expressing CD4∷mIFP, Par3∷YFP and constitutively active FGFR, *λbtl*, under *btl-gal4*. (C’-D’) CD4∷mIFP. (C’’-E’’) Manual tracings of the cell contours (dark gray), subcellular tube (black), endosomes (red) and dorsal branch cells (light gray).

**Figure S2.**
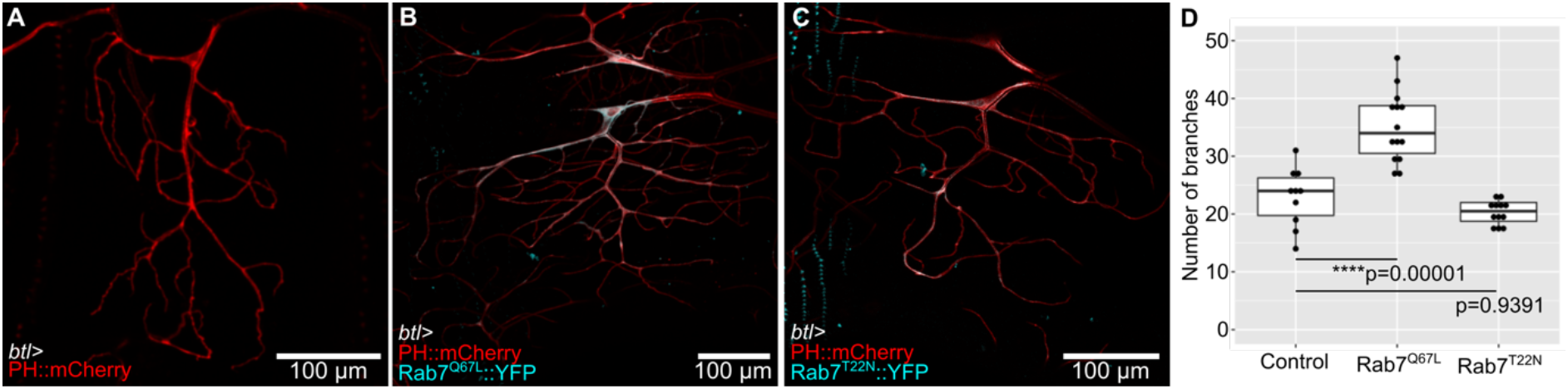
Loss- and gain of Rab7 function and its role in larval terminal cell branching. Larval terminal cells expressing PH∷mCherry under *btl-gal4*. (A) Control, TM6BTb cell, (B) sibling cell additionally expressing Rab7^Q67L^∷YFP, (C) cell expressing Rab7^T22N^∷YFP. Significance was assayed using one-way ANOVA and Tukey correction. Number of cells analyzed: control, n=10; Rab7^Q67L^∷YFP, n=14, Rab7^T22N^∷YFP, n=12.

**Figure S3.**
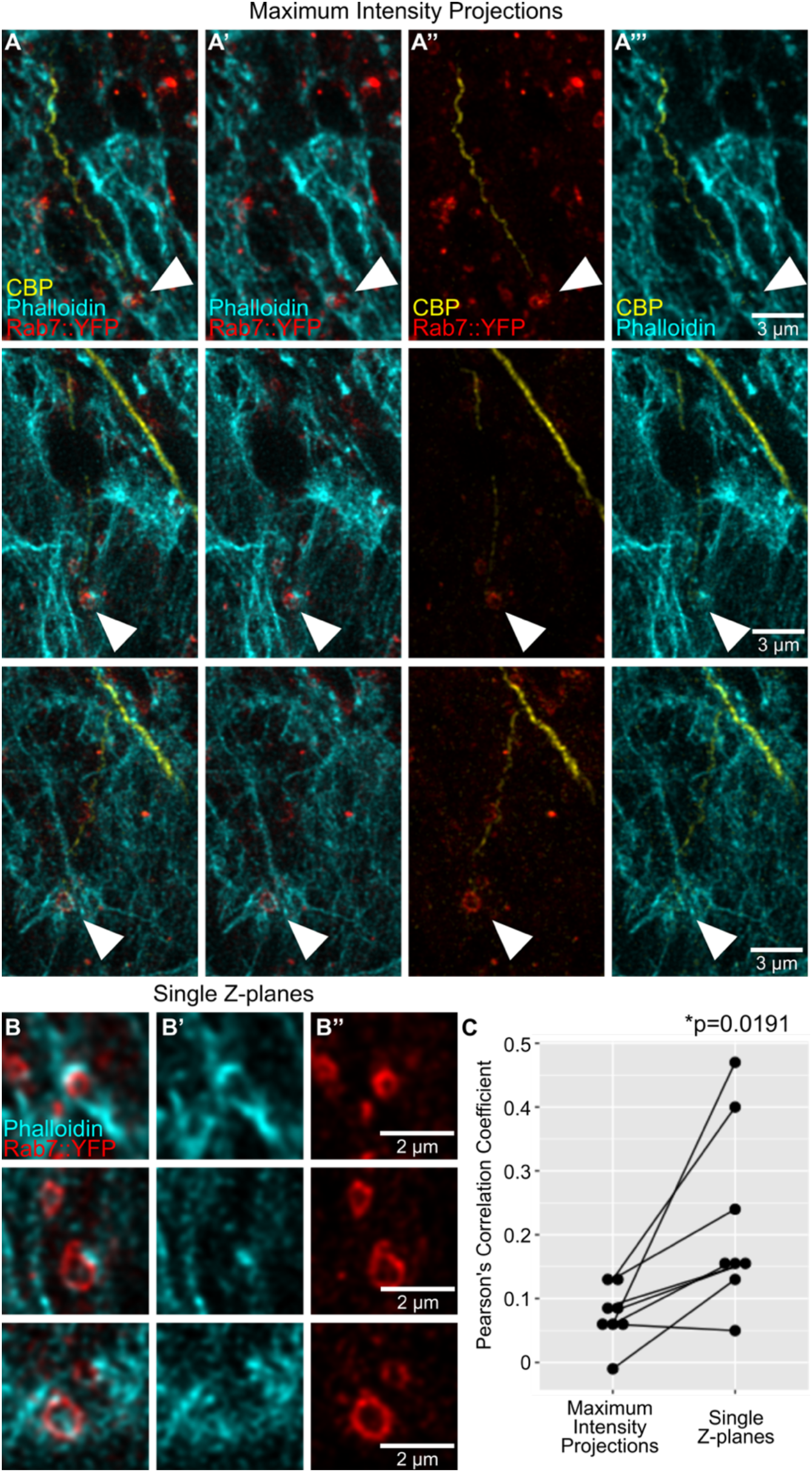
Actin enrichment around late endosomes. (A-B) Terminal cells expressing Rab7∷YFP and stained with phalloidin and a chitin binding protein (CBP) fused to Alexa647. (A-A’’’) Channels shown in pair-wise combinations. Arrowheads point to the contact between Rab7∷YFP rings, actin and the tip of the subcellular tube. (B) Close-ups of the same cells as in (A), but shown as single Z-planes. Colors are the same as in (A). (C) Correlation between pixel fluorescence intensities in the phalloidin and YRab7 channels, in maximum intensity projections (as shown in panels A) vs single Z-planes (as shown in panels B). Significance was assessed using two-tailed paired t-test, n=8.

**Figure S4.**
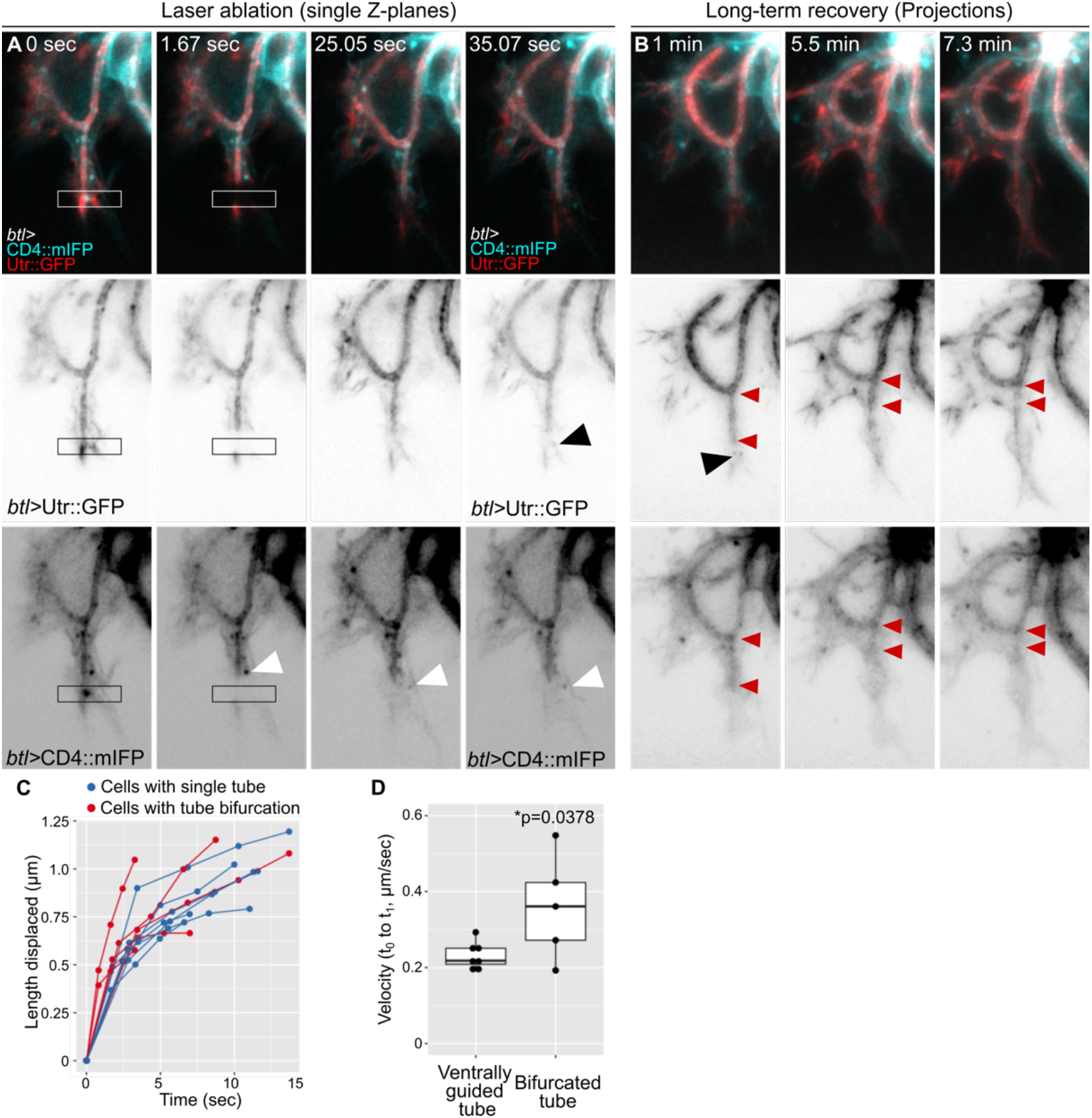
Infrared laser ablation at the tip of terminal cells with bifurcated tubes. (A-B) Terminal cell expressing CD4∷mIFP and Utr∷GFP under *btl-gal4*. (A) Cell immediately before laser cut. The white box marks the laser-ablated region. (B) Long-term recovery, z-projections. (C-D) Quantification of the recoil as determined by length displacement over time, using manual tracking. (C) Time course, (D) Initial velocity. Number of cells analysed: cells with properly guided tube, n=7; cells with misguided tube, n=5. Significance was assessed using t-test.

**Figure S5.**
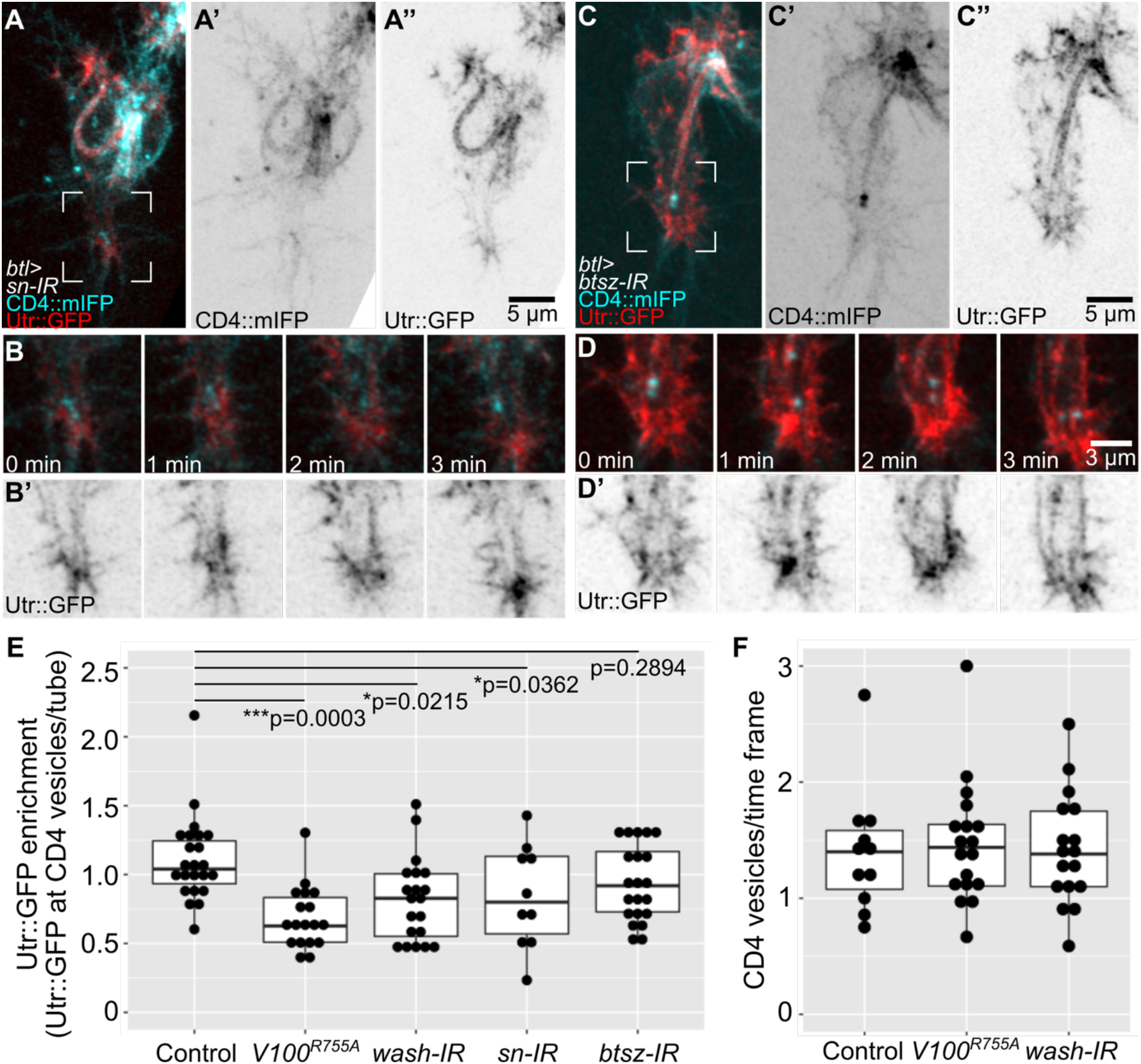
Actin organization in cells with defective actin-binding proteins. (A-D) Cells expressing CD4∷mIFP and Utr∷GFP under *btl-gal4*. (A-B) *sn* knockdown, (C-D) *btsz* knockdown. Regions marked in A and C are magnified and shown over four time points in B and D, respectively. (E) Mean Utr∷GFP fluorescence intensity in CD4 vesicles over time. Values were normalized to fluorescence intensity at the tube to adjust for variations in the expression of the Utr∷GFP reporter. Number of vesicles analyzed: control, n=23 from 9 cells; Vha100^R755A^, n=19 from 8 cells; *wash-IR*, n=20 from 9 cells; *sn-IR,* n=10 from 5 cells; *btsz-IR*, n=20 from 8 cells. Values at the top are significance between genotypes, assessed by ANOVA and Tukey correction for multiple comparisons. (F) Mean number of CD4 vesicles at the tip of the cell per timepoint (4-28 timepoints per cell). Control cells, n=11; *Vha100^R755A^*, n=18; *wash* RNAi, n=17.

## Supplemental Movies

Movie S1. Terminal cell development in *shrb* mutants and control siblings.

Movie S2. Rab7 mislocalization using the GrabFP system.

Movie S3. High temporal resolution imaging of CD4∷mIFP and Utrophin∷GFP distribution during terminal cell development.

Movie S4. Laser cuts at the growing tip of terminal cells.

Movie S5. Distribution of CD4∷mIFP and Utr∷GFP during terminal cell development in control cells, in cells expressing *Vha10^0R755A^* and in cells expressing *wash-IR.*

## References

Alekhina, O., E. Burstein, and D.D. Billadeau. 2017. Cellular functions of WASP family proteins at a glance. J. Cell Sci. 130:2235–2241. doi:10.1242/jcs.199570.

Best, B.T. 2019. Single-cell branching morphogenesis in the Drosophila trachea. Dev Biol. 451:5–15. doi:10.1016/j.ydbio.2018.12.001.

Boda, A., P. Lőrincz, S. Takáts, T. Csizmadia, S. Tóth, A.L. Kovács, and G. Juhász. 2019. Drosophila Arl8 is a general positive regulator of lysosomal fusion events. Biochim. Biophys. Acta - Mol. Cell Res. 1866:533–544. doi:10.1016/j.bbamcr.2018.12.011.

Bun, P., S. Dmitrieff, J.M. Belmonte, F.J. Nédélec, and P. Lénárt. 2018. A disassembly-driven mechanism explains F-actin-mediated chromosome transport in starfish oocytes. Elife. 7. doi:10.7554/eLife.31469.

Chanut-Delalande, H., A.C. Jung, M.M. Baer, L. Lin, F. Payre, and M. Affolter. 2010. The Hrs/Stam complex acts as a positive and negative regulator of RTK signaling during Drosophila development. PLoS One. 5:e10245. doi:10.1371/journal.pone.0010245.

Chanut-Delalande, H., A.C. Jung, L. Lin, M.M. Baer, A. Bilstein, C. Cabernard, M. Leptin, and M. Affolter. 2007. A genetic mosaic analysis with a repressible cell marker screen to identify genes involved in tracheal cell migration during Drosophila air sac morphogenesis. Genetics. 176:2177–2187. doi:10.1534/genetics.107.073890.

Cherry, S., E.J. Jin, M.N. Özel, Z. Lu, E. Agi, D. Wang, W.H. Jung, D. Epstein, I.A. Meinertzhagen, C.C. Chan, and P. Robin Hiesinger. 2013. Charcot-Marie-Tooth 2B mutations in rab7 cause dosage-dependent neurodegeneration due to partial loss of function. Elife. 2013. doi:10.7554/eLife.01064.001.

Choi, W., K.-C. Jung, K.S. Nelson, M. a Bhat, G.J. Beitel, M. Peifer, and A.S. Fanning. 2011. The single Drosophila ZO-1 protein Polychaetoid regulates embryonic morphogenesis in coordination with Canoe/afadin and Enabled. Mol. Biol. Cell. 22:2010–30. doi:10.1091/mbc.E10-12-1014.

Derivery, E., C. Sousa, J.J. Gautier, B. Lombard, D. Loew, and A. Gautreau. 2009. The Arp2/3 Activator WASH Controls the Fission of Endosomes through a Large Multiprotein Complex. Dev. Cell. 17:712–723. doi:10.1016/j.devcel.2009.09.010.

Dogterom, M., and G.H. Koenderink. 2019. Actin–microtubule crosstalk in cell biology. Nat. Rev. Mol. Cell Biol. 20:38–54. doi:10.1038/s41580-018-0067-1.

Dong, B., E. Hannezo, and S. Hayashi. 2014. Balance between apical membrane growth and luminal matrix resistance determines epithelial tubule shape. Cell Rep. 7:941–950. doi:10.1016/j.celrep.2014.03.066.

Dong, B., K. Kakihara, T. Otani, H. Wada, and S. Hayashi. 2013. Rab9 and retromer regulate retrograde trafficking of luminal protein required for epithelial tube length control. Nat. Commun. 4:1312–1358. doi:10.1038/ncomms2347.

Dunst, S., T. Kazimiers, F. von Zadow, H. Jambor, A. Sagner, B. Brankatschk, A. Mahmoud, S. Spannl, P. Tomancak, S. Eaton, and M. Brankatschk. 2015. Endogenously tagged rab proteins: a resource to study membrane trafficking in Drosophila. Dev Cell. 33:351–365. doi:10.1016/j.devcel.2015.03.022.

Francis, D., and A.S. Ghabrial. 2015. Compensatory branching morphogenesis of stalk cells in the Drosophila trachea. Development. 142:2048–2057. doi:10.1242/dev.119602.

Gervais, L., and J. Casanova. 2010. In vivo coupling of cell elongation and lumen formation in a single cell. Curr Biol. 20:359–366. doi:10.1016/j.cub.2009.12.043.

Gilmour, D., M. Rembold, and M. Leptin. 2017. From morphogen to morphogenesis and back. Nature. 541:311–320. doi:10.1038/nature21348.

Gomez, T.S., J.A. Gorman, A.A.M. de Narvajas, A.O. Koenig, and D.D. Billadeau. 2012. Trafficking defects in WASH-knockout fibroblasts originate from collapsed endosomal and lysosomal networks. Mol. Biol. Cell. 23:3215–3228. doi:10.1091/mbc.e12-02-0101.

Goodwin, K., S.J. Ellis, E. Lostchuck, T. Zulueta-Coarasa, R. Fernandez-Gonzalez, and G. Tanentzapf. 2016. Basal Cell-Extracellular Matrix Adhesion Regulates Force Transmission during Tissue Morphogenesis. Dev. Cell. 39:611–625. doi:10.1016/j.devcel.2016.11.003.

Harmansa, S., I. Alborelli, D. Bieli, E. Caussinus, and M. Affolter. 2017. A nanobody-based toolset to investigate the role of protein localization and dispersal in Drosophila. Elife. 6. doi:10.7554/eLife.22549.

Hayashi, S., and T. Kondo. 2018. Development and function of the drosophila tracheal system. Genetics. 209:367–380. doi:10.1534/genetics.117.300167.

Hiesinger, P.R., A. Fayyazuddin, S.Q. Mehta, T. Rosenmund, K.L. Schulze, R.G. Zhai, P. Verstreken, Y. Cao, Y. Zhou, J. Kunz, and H.J. Bellen. 2005. The v-ATPase V0 subunit a1 is required for a late step in synaptic vesicle exocytosis in Drosophila. Cell. 121:607–620. doi:10.1016/j.cell.2005.03.012.

JayaNandanan, N., R. Mathew, and M. Leptin. 2014. Guidance of subcellular tubulogenesis by actin under the control of a synaptotagmin-like protein and Moesin. Nat Commun. 5:3036. doi:10.1038/ncomms4036.

Kasza, K.E., S. Supriyatno, and J.A. Zallen. 2019. Cellular defects resulting from disease-related myosin II mutations in Drosophila. Proc. Natl. Acad. Sci. U. S. A. 116:22205–22211. doi:10.1073/pnas.1909227116.

Lebreton, G., and J. Casanova. 2016. Ligand-binding and constitutive FGF receptors in single Drosophila tracheal cells: Implications for the role of FGF in collective migration. Dev Dyn. 245:372–378. doi:10.1002/dvdy.24345.

Lee, T., N. Hacohen, M. Krasnow, and D.J. Montell. 1996. Regulated breathless receptor tyrosine kinase activity required to pattern cell migration and branching in the Drosophila tracheal system. Genes Dev. 10:2912–2921. doi:10.1101/gad.10.22.2912.

Levi, B.P., A.S. Ghabrial, and M.A. Krasnow. 2006. Drosophila talin and integrin genes are required for maintenance of tracheal terminal branches and luminal organization. Development. 133:2383–2393. doi:10.1242/dev.02404.

MacDonald, E., L. Brown, A. Selvais, H. Liu, T. Waring, D. Newman, J. Bithell, D. Grimes, S. Urbe, M.J. Clague, and T. Zech. 2018. HRS-WASH axis governs actin-mediated endosomal recycling and cell invasion. J Cell Biol. 217:2549–2564. doi:10.1083/jcb.201710051.

Mathew, R., L.D. Rios-Barrera, P. Machado, Y. Schwab, and M. Leptin. 2020. Transcytosis via the late endocytic pathway as a cell morphogenetic mechanism. EMBO J. 2020.01.16.909200. doi:10.15252/embj.2020105332.

Nagel, B.M., M. Bechtold, L.G. Rodriguez, and S. Bogdan. 2017. Drosophila WASH is required for integrin-mediated cell adhesion, cell motility and lysosomal neutralization. J. Cell Sci. 130:344–359. doi:10.1242/jcs.193086.

Nikolova, L.S., and M.M. Metzstein. 2015. Intracellular lumen formation in Drosophila proceeds via a novel subcellular compartment. Development. 142:3964–3973. doi:10.1242/dev.127902.

Ochoa-Espinosa, A., S. Harmansa, E. Caussinus, and M. Affolter. 2017. Myosin II is not required for Drosophila tracheal branch elongation and cell intercalation. J. Cell Sci. 130:2961–2968. doi:10.1242/dev.148940.

Okenve-Ramos, P., and M. Llimargas. 2014. Fascin links Btl/FGFR signalling to the actin cytoskeleton during Drosophila tracheal morphogenesis. Development. 141:929–939. doi:10.1242/dev.103218.

Oshima, K., M. Takeda, E. Kuranaga, R. Ueda, T. Aigaki, M. Miura, and S. Hayashi. 2006. IKK epsilon regulates F actin assembly and interacts with Drosophila IAP1 in cellular morphogenesis. Curr Biol. 16:1531–1537. doi:10.1016/j.cub.2006.06.032.

Palamidessi, A., E. Frittoli, M. Garré, M. Faretta, M. Mione, I. Testa, A. Diaspro, L. Lanzetti, G. Scita, and P.P. Di Fiore. 2008. Endocytic Trafficking of Rac Is Required for the Spatial Restriction of Signaling in Cell Migration. Cell. 134:135–147. doi:10.1016/j.cell.2008.05.034.

Piacentino, M.L., Y. Li, and M.E. Bronner. 2020. Epithelial-to-mesenchymal transition and different migration strategies as viewed from the neural crest. Curr. Opin. Cell Biol. 66:43–50. doi:10.1016/j.ceb.2020.05.001.

Ricolo, D., and S.J. Araujo. 2020. Coordinated crosstalk between microtubules and actin by a spectraplakin regulates lumen formation and branching. Elife. 9:2020.05.09.085753. doi:10.7554/eLife.61111.

Ricolo, D., M. Deligiannaki, J. Casanova, and S.J. Araujo. 2016. Centrosome Amplification Increases Single-Cell Branching in Post-mitotic Cells. Curr Biol. 26:2805–2813. doi:10.1016/j.cub.2016.08.020.

Rosa-Ferreira, C., S.T. Sweeney, and S. Munro. 2018. The small G protein Arl8 contributes to lysosomal function and long-range axonal transport in Drosophila. Biol. Open. 7. doi:10.1242/bio.035964.

Rothwell, W.F., and W. Sullivan. 2007. Fixation of Drosophila Embryos. Cold Spring Harb. Protoc. 2007:pdb.prot4827–pdb.prot4827. doi:10.1101/pdb.prot4827.

Rottner, K., J. Faix, S. Bogdan, S. Linder, and E. Kerkhoff. 2017. Actin assembly mechanisms at a glance. J. Cell Sci. 130:3427–3435. doi:10.1242/jcs.206433.

Schiefermeier, N., J.M. Scheffler, M.E.G. de Araujo, T. Stasyk, T. Yordanov, H.L. Ebner, M. Offterdinger, S. Munck, M.W. Hess, S.A. Wickström, A. Lange, W. Wunderlich, R. Fässler, D. Teis, and L.A. Huber. 2014. The late endosomal p14-MP1 (LAMTOR2/3) complex regulates focal adhesion dynamics during cell migration. J. Cell Biol. 205:525–540. doi:10.1083/jcb.201310043.

Schottenfeld-Roames, J., J.B. Rosa, and A.S. Ghabrial. 2014. Seamless tube shape is constrained by endocytosis-dependent regulation of active Moesin. Curr Biol. 24:1756–1764. doi:10.1016/j.cub.2014.06.029.

Shaye, D.D., and I. Greenwald. 2015. The disease-associated formin INF2/EXC-6 organizes lumen and cell outgrowth during tubulogenesis by regulating F-actin and microtubule cytoskeletons. Dev. Cell. 32:743–755. doi:10.1016/j.devcel.2015.01.009.

Shcherbata, H.R., A.S. Yatsenko, L. Patterson, V.D. Sood, U. Nudel, D. Yaffe, D. Baker, and H. Ruohola-Baker. 2007. Dissecting muscle and neuronal disorders in a Drosophila model of muscular dystrophy. EMBO J. 26:481–493. doi:10.1038/sj.emboj.7601503.

Shiga, Y., M. TanakaMatakatsu, S. Hayashi, M. Tanaka-Matakatsu, and S. Hayashi. 1996. A nuclear GFP beta-galactosidase fusion protein as a marker for morphogenesis in living Drosophila. Dev. Growth Differ. 38:99–106. doi:10.1046/j.1440-169X.1996.00012.x.

Sigurbjörnsdóttir, S., R. Mathew, and M. Leptin. 2014. Molecular mechanisms of de novo lumen formation. Nat Rev Mol Cell Biol. 15:665–676. doi:10.1038/nrm3871.

Siller, K.H., C. Cabernard, and C.Q. Doe. 2006. The NuMA-related Mud protein binds Pins and regulates spindle orientation in Drosophila neuroblasts. Nat Cell Biol. 8:594–600. doi:10.1038/ncb1412.

Sitarska, E., and A. Diz-Muñoz. 2020. Pay attention to membrane tension: Mechanobiology of the cell surface. Curr. Opin. Cell Biol. 66:11–18. doi:10.1016/j.ceb.2020.04.001.

Spracklen, A.J., T.N. Fagan, K.E. Lovander, and T.L. Tootle. 2014. The pros and cons of common actin labeling tools for visualizing actin dynamics during Drosophila oogenesis. Dev. Biol. 393:209–226. doi:10.1016/j.ydbio.2014.06.022.

Sundaram, M. V, and J.D. Cohen. 2017. Time to make the doughnuts: Building and shaping seamless tubes. Semin Cell Dev Biol. 67:123–131. doi:10.1016/j.semcdb.2016.05.006.

Tsarouhas, V., D. Liu, G. Tsikala, A. Fedoseienko, K. Zinn, R. Matsuda, D.D. Billadeau, and C. Samakovlis. 2019. WASH phosphorylation balances endosomal versus cortical actin network integrities during epithelial morphogenesis. Nat Commun. 10:2193. doi:10.1038/s41467-019-10229-6.

Tseng, Q., E. Duchemin-Pelletier, A. Deshiere, M. Balland, H. Guilloud, O. Filhol, and M. Theŕy. 2012. Spatial organization of the extracellular matrix regulates cell-cell junction positioning. Proc. Natl. Acad. Sci. U. S. A. 109:1506–1511. doi:10.1073/pnas.1106377109.

Walton, K.D., A.M. Freddo, S. Wang, and D.L. Gumucio. 2016. Generation of intestinal surface: An absorbing tale. Dev. 143:2261–2272. doi:10.1242/dev.135400.

Williamson, W.R., D. Wang, A.S. Haberman, and P.R. Hiesinger. 2010. A dual function of V0-ATPase a1 provides an endolysosomal degradation mechanism in Drosophila melanogaster photoreceptors. J Cell Biol. 189:885–899. doi:10.1083/jcb.201003062.

Zhang, J., K.L. Schulze, P. Robin Hiesinger, K. Suyama, S. Wang, M. Fish, M. Acar, R.A. Hoskins, H.J. Bellen, and M.P. Scott. 2007. Thirty-one flavors of Drosophila Rab proteins. Genetics. 176:1307–1322. doi:10.1534/genetics.106.066761.

